# Natural Microbial Enrichment Modulates Microglial States and Transcriptional Programs Relevant to Alzheimer’s Disease

**DOI:** 10.64898/2026.02.28.708710

**Authors:** Mohamed Lala Bouali, Amir Mohamed Kezai, Marc Bazin, Papa Yaya Badiane, Nasim Eskandari, Véronique Lévesque, Justin Robillard, Steeve D. Côté, Denis Soulet, Cyntia Tremblay, Frédéric Calon, Françoise Morin, Luc Vallières, Sébastien S. Hébert

## Abstract

The diversity of environmental microbial exposure is a key driver of immune maturation and host defense; however, its impact on brain immunity and neurodegenerative diseases remains poorly documented. Here, we show that controlled indoor rewilding, by introducing a natural farm-like environment into laboratory housing, reshapes peripheral and central nervous system (CNS) immune networks in wild-type (WT) and 5xFAD mice, a model of Alzheimer’s disease (AD). Compared with traditional specific pathogen-free (SPF) housing, rewilded mice exhibited systemic shifts toward mature immune phenotypes, including increases in effector and memory B and T cells, expansion of plasma cell populations, and alterations in immunoglobulin isotypes. In the brain, indoor rewilding recalibrated microglial activation of SPF-5xFAD mice, attenuating pro-inflammatory transcriptional programs while enhancing homeostatic, complement, and phagocytic signatures. A strong transcriptional convergence was observed between rewilded and wild mice, with rewilded 5xFAD mice exhibiting greater similarity to human AD transcriptional profiles. Morphological and histochemical analyses confirmed that rewilded microglia adopt metabolically adaptable, homeostatic states that influence amyloid-β plaque binding and clearance. Collectively, these findings suggest that microbial diversity through “dirty” mouse modeling could enhance the translational relevance of neuroimmunology and neurodegenerative disease research.

## INTRODUCTION

The peripheral and CNS immune systems interact in a continuous and bidirectional manner to orchestrate host defense, maintain tissue homeostasis, and resolve inflammation^1–3^. Microglia, the primary resident immune cells of the brain, integrate signals from neurons, astrocytes, endothelial cells, and perivascular macrophages, while remaining responsive to cues derived from the peripheral immune system, such as lymphocytes. Through this extensive bidirectional crosstalk, microglia regulate processes such as cytokine production, antigen presentation, phagocytosis, and synaptic pruning^1,3,4^. Although recent studies have demonstrated that systemic triggers can influence immune dynamics within the brain, our understanding of how real-world immune challenges impact microglial activity remains limited. Indeed, standard laboratory murine models raised under hygienic specific pathogen-free (SPF) conditions possess naïve and underdeveloped immune systems, characterized by low populations of effector and memory T and B cells, reduced immunoglobulin class switching, altered phenotypes of innate immune cells, and diminished or skewed inflammatory responses^5–7^. This limited exposure to natural microbes and complex antigens likely affects microglial states and functions.

The growing field of eco-immunology has shown that restoring natural antigenic diversity can recalibrate immune priming and maturation, thereby improving physiological and disease modeling in laboratory mice^8,9^. Strategies include wildling models^10,11^, naturalization or feralization methods^5,12–15^, and rewilding paradigms^16,17^, all of which have shown that exposure to natural microorganisms and fomites promotes genuine immune development, comparable to that seen in wild-caught mice and adult humans^10,18–20^. Positive outcomes include increased resistance to pathogens, human-like immunization responses, and adapted inflammatory reactions across different organs such as the blood, spleen, gut, skin, and lungs.

Although these “dirty” modelling approaches reveal the critical role of antigenic diversity and complexity in shaping peripheral immunity, their impact on CNS-resident immune cells, particularly microglia, remains poorly characterized. One report has described how rewilding affects microglial activity in the context of fungal infection^21^. Whether environmentally driven immune maturation modulates microglial states in contexts of chronic neuroinflammation or neurodegenerative disease, such as Alzheimer’s disease (AD), remains unknown.

In AD, microglia transition from homeostatic surveillance states to disease-associated phenotypes (e.g., disease-associated microglia, DAM), characterized by altered gene expression, morphology, and function^22–24^. These activated microglial states play complex roles in the metabolism (e.g., phagocytosis, compaction, containment, and clearance) of pathological amyloid-β (Aβ) plaques and neuroinflammation^25–27^. Although recent studies indicate that peripheral stimuli can influence CNS immune responses and disease outcomes in SPF AD mice^28–30^, it remains uncertain whether their immunological naïveté affects microglial transcriptional states and functions. This issue is vital for understanding the similarities and differences between murine and human AD microglial cells and, by extension, other immune cell responses^22,31–36^.

In the current work, we employed a controlled indoor rewilding paradigm^37^ to examine how a semi-natural farm-like environment influences peripheral and cerebral immune composition in wildtype (WT) and 5xFAD mice, a well-established transgenic model of Aβ pathology used in AD immunological research^38–43^. Our findings demonstrate that natural environmental exposure induces extensive immune maturation and remodeling, redirects brain transcriptional signatures towards those observed in human AD profiles, and reshapes Aβ-associated microglial responses. These findings position dirty mouse modeling as a potent driver of authentic neurodegenerative disease biology, offering a pathway toward more predictive, realistic, and human-relevant preclinical models.

## RESULTS

### Indoor rewilding recalibrates systemic leukocyte composition in WT and 5xFAD mice

To examine how indoor rewilding impacts systemic immunity in an AD model, two-month-old SPF-born WT and 5xFAD female mice were randomly assigned to four experimental groups (**Figure 1B**). SPF cohorts (SPF-WT and SPF-5xFAD, N = 14 per genotype) were maintained in standard SPF animal facility conditions (IVCs). In parallel, rewilded cohorts (Rewild-WT and Rewild-5xFAD, N = 14 per genotype) were released into semi-natural enclosures (**Figure 1A**) for three months, a time period consistent with prior rewilding or naturalization studies^12,16,44^. To mimic natural *Mus musculus* habitats, the enclosures were enriched with a diverse array of organic and living biological elements, including farmland soil, plants, and arthropods, as described before^37^. No external pathogens are introduced using this method^37^, and no mortality or health issues were observed in the current study. All animals were euthanized at five months of age (**Figure 1C**), corresponding to an early-to-mid stage of AD pathology in 5xFAD mice marked by Aβ plaque deposition, microglial activation, and neuroinflammation, yet preceding neurodegeneration or cognitive and motor decline^38,40,45^. Consistent with our previous findings^37^, rewilded 5xFAD mice were, on average, 2.5 grams leaner than their SPF counterparts (not shown), likely reflecting increased physical activity and metabolic expenditure.

**Figure 1.**
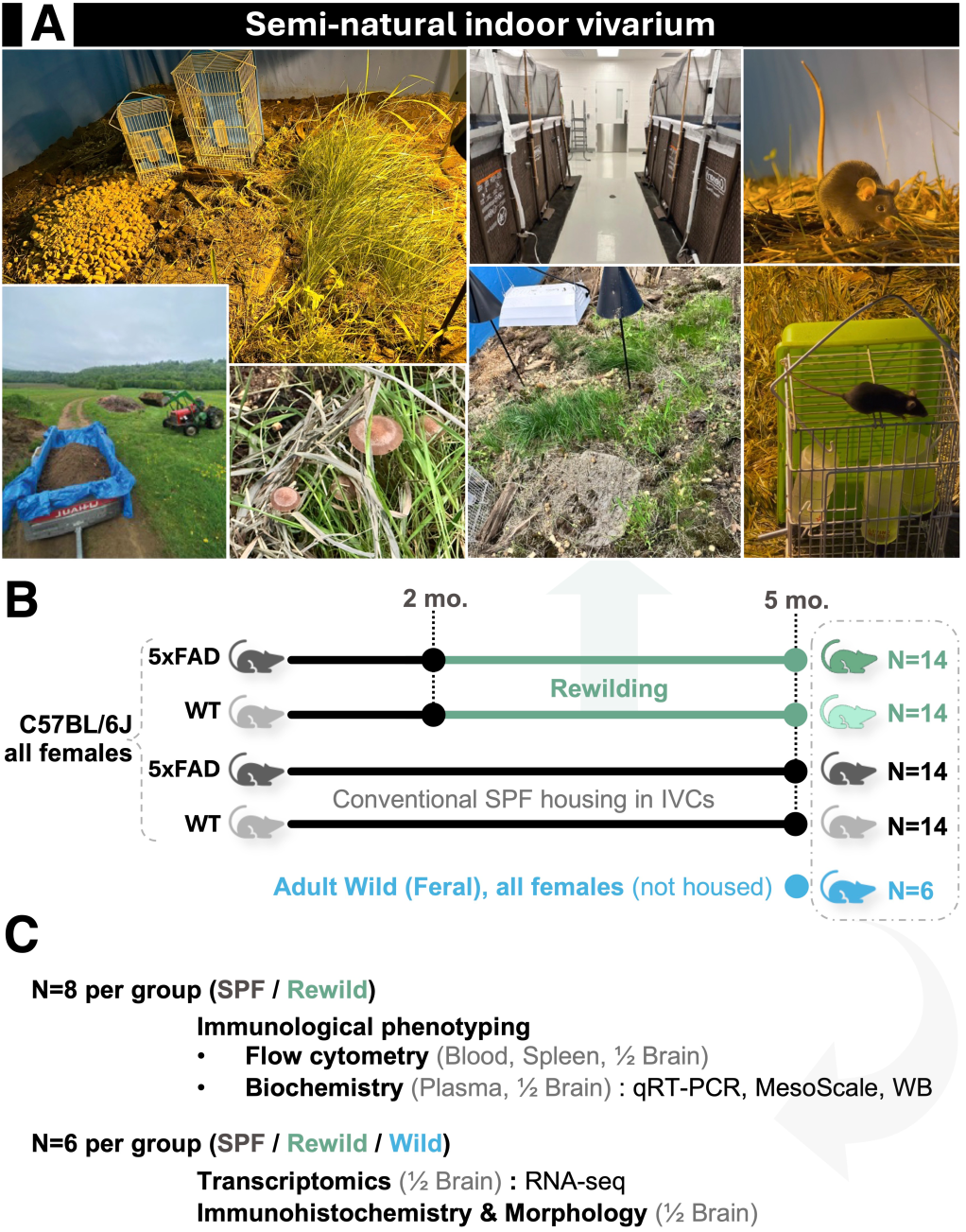
Experimental design, housing conditions, and cohort allocation across assays. (A) Representative photographs of the semi-natural indoor vivarium used for the rewilding exposure. A detailed description of the vivarium platform (structure, substrates, enrichment elements, and environmental features) was reported previously^37^. (B) Two-month-old female SPF 5xFAD mice and their age-matched female WT littermates (SPF-WT) mice underwent routine health assessment and baseline body-weight measurement, then were randomly assigned to standard SPF housing in individually ventilated cages (IVCs) or rewilding housing in the indoor vivarium (N=14 per group): WT-SPF, WT-Rewild, 5xFAD-SPF, and 5xFAD-Rewild. WT and 5xFAD mice were housed in two separate, strictly identical vivariums located side-by-side in the same room to avoid mixing of visually indistinguishable mice on the same C57BL/6J background. Rewilding was conducted for 3 months (2 to 5 months of age). At study endpoint, rewilded mice were captured using Havahart-type live traps and processed for downstream assays. (C) For immunophenotyping and biochemistry, N=8 per group (SPF vs Rewild) were used for flow cytometry (blood, spleen, and right brain hemisphere) and biochemical assays (plasma and left brain hemisphere; qRT-PCR, Meso Scale Discovery (MSD) multiplex immunoassays, and immunoblotting). For transcriptomics and histology, N=6 per group were used for brain RNA sequencing (left hemisphere) and immunohistochemistry/morphology (right hemisphere, fixed). An additional group of adult Wild (Feral) female mice (N=6), captured outdoors and euthanized immediately (not housed under SPF or rewilding conditions), was included only for brain transcriptomic profiling (RNA-seq). Flow cytometry data are shown in Figures 2-7, biochemical analyses in Figures 7-8,10, immunohistochemistry/morphology in Figure 8, and transcriptomic analyses in Figures 9-11.

Rewilding increased the absolute number of CD45⁺ leukocytes in both spleen and blood regardless of genotype (**Figure 2A,C**). However, the proportion of CD45⁺ cells among live events decreased under rewilding in both tissues, consistent with a housing-dependent shift in the overall composition of acquired suspensions. The magnitude of these effects differed by tissue: the increase in CD45⁺ cell number was more pronounced in blood than in spleen, whereas the reduction in absolute CD45⁺ cell proportion was larger in spleen than in blood. In 5xFAD mice, this reduction was consistent with a proportionally larger increase in non-CD45⁺ live events under rewilding (**Figure 2A,C**).

**Figure 2.**
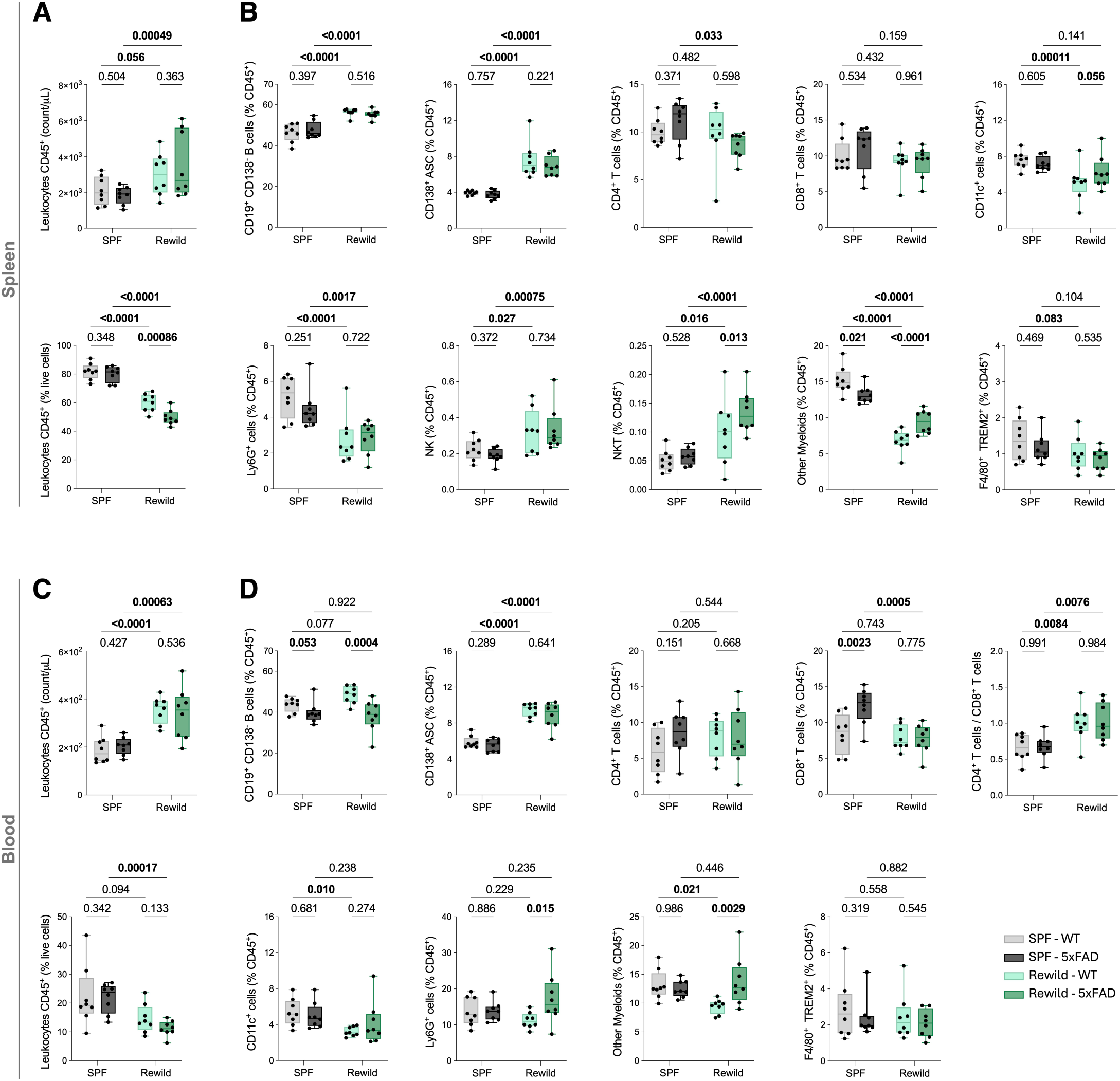
Rewilding reshapes systemic leukocyte composition in spleen and blood. Flow cytometry quantification of leukocytes in spleen (A-B) and blood (C-D) from female mice housed under SPF conditions or in the indoor vivarium (Rewild): SPF-WT, SPF-5xFAD, Rewild-WT, and Rewild-5xFAD (N = 8 per group). Gating strategies are provided in Supplementary Figure 1. (A, C) Total leukocytes are shown as numbers (volumetric counts; CD45⁺ events per µL of acquired sample; acquisition target ∼100µL per sample) and as proportions (CD45⁺, % of live cells). (B, D) proportions of major leukocyte subsets are expressed as % of CD45⁺. In spleen, subsets include CD138⁻CD19⁺ B cells, CD138⁺ antibody-secreting cells (ASCs), CD4⁺ T cells, CD8⁺ T cells, NK, NKT, Ly6G⁺ cells, CD11c⁺ cells, F4/80⁺TREM2⁺ macrophages, and other myeloids (including macrophage and monocyte populations). In blood, only subsets detectable in circulation are shown, and the CD4⁺/CD8⁺ T-cells ratio is included. Data are plotted as individual values overlaid on box-and-whisker plots (box: 25^th^-75^th^ percentile; center line: median; whiskers: min-max). Statistical analyses were performed in R, as described in Methods. Numbers for leukocyte subsets (counts/µL) are provided in Supplementary Figure 2.

In the spleen, rewilding strongly expanded the B cell compartment. Both CD19⁺CD138⁻ B cells and CD138⁺ antibody-secreting cells (ASCs) increased under rewilding, not only as a fraction of CD45⁺ leukocytes (**Figure 2B**) but also in absolute number (**Supplementary Figure 2A**), indicating that these changes reflect increased B-lineage cellular output rather than a purely compositional redistribution. Rewilding also increased the number (**Figure 2B**) and proportion (**Supplementary Figure 2A**) of innate lymphoid cells (NK and NKT) in spleen, consistent with coordinated expansion of these compartments.

In contrast, several myeloid-associated changes were primarily compositional within the spleen. Ly6G⁺ cells decreased in proportion among leukocytes (**Figure 2B**) while absolute Ly6G⁺ cell number remained largely unchanged (**Supplementary Figure 2A**), consistent with a relative redistribution within the splenic leukocyte pool.

Similarly, the proportion of other myeloid cells decreased substantially in the spleen (**Figure 2B**) without a corresponding decrease in absolute number (**Supplementary Figure 2A**), indicating that the reduced frequency largely reflects remodeling of the CD45⁺ compartment rather than a net loss of these events. Notably, this compartment displayed an environment-dependent genotype pattern that reversed across housing conditions (SPF-WT > SPF-5xFAD; Rewild-WT < Rewild-5xFAD) (**Figure 2B**).

CD11c⁺ cells showed a decrease that was more evident in WT under rewilding (**Figure 2B**), with only modest changes in abundance (**Supplementary Figure 2A**). F4/80⁺TREM2⁺ macrophages showed a modest downward trend in frequency with limited change in abundance.

In the blood, rewilding substantially increased the number and proportion of ASCs (**Figure 2D** and **Supplementary Figure 2B**). By contrast, CD19⁺CD138⁻ B cell number was stable across environments, while absolute B cell number increased under rewilding (**Supplementary Figure 2B**). Importantly, CD19⁺CD138⁻ B cells showed genotype-dependent changes in number and proportion that increased under rewilding. In addition, 5xFAD mice had significantly higher number and proportion of CD8^+^ cytotoxic T cells in SPF conditions, which was recalibrated to WT level under rewilding (**Figure 2D**). Furthermore, the CD4⁺/CD8⁺ T cell ratio increased under rewilding conditions (**Figure 2D**), consistent with a more pronounced regulatory tone within the blood compartment.

Blood myeloid populations showed distinct, genotype-dependent changes. Ly6G⁺ cells increased in number under rewilding (**Supplementary Figure 2B**) along with a significant proportion increase in 5xFAD (**Figure 2D**). For other myeloid cells, number and proportion diverged across genotype. Finally, CD11c^+^ cells showed a similar pattern as in spleen with significant decrease in WT under rewilding (**Figure 2D**). Taken together, these data show that indoor rewilding broadly remodels systemic immunity in both WT and 5xFAD mice, characterized by increased leukocyte abundance alongside pronounced shifts in leukocyte composition.

### Rewilding increases B cell maturation and activation

Among the splenic CD19⁺CD138⁻ B cell population, the proportion of cells expressing multiple activation-associated markers (CD38, CD44, CD25) was increased by rewilding (**Figure 3A**). However, the Rewild-WT group showed a decreased proportion of CD86⁺ B cells. In contrast, the 5xFAD group showed an increased proportion of CD335⁺ B cells. Consistent with a differentiation shift, rewilding reduced the proportion of naïve B cells (CD38⁺IgD⁺) in spleen (**Figure 3B**; environment effect shown in the adjacent interaction plot), while also decreasing germinal center-like B cells (CD38⁻IgD⁻; **Figure 3C**; environment effect), but increasing memory-like B cells (CD38⁺IgD⁻) both in proportion and number (**Figure 3D**; **Supplementary Figure 3A**; environment effect). Notably, this latter subset showed marked population heterogeneity that was partitioned based on CD38 intensity and FSC-H (**Figure 3D**, right plot), motivating deeper quantification in **Supplementary Figure 3**. Phenotypically, these CD38/FSC-defined subsets differed in their expression of CD19, CD38, CD44, and CD86. Genotype differences were consistent across environmental conditions (**Supplementary Figure 3B-D**).

**Figure 3.**
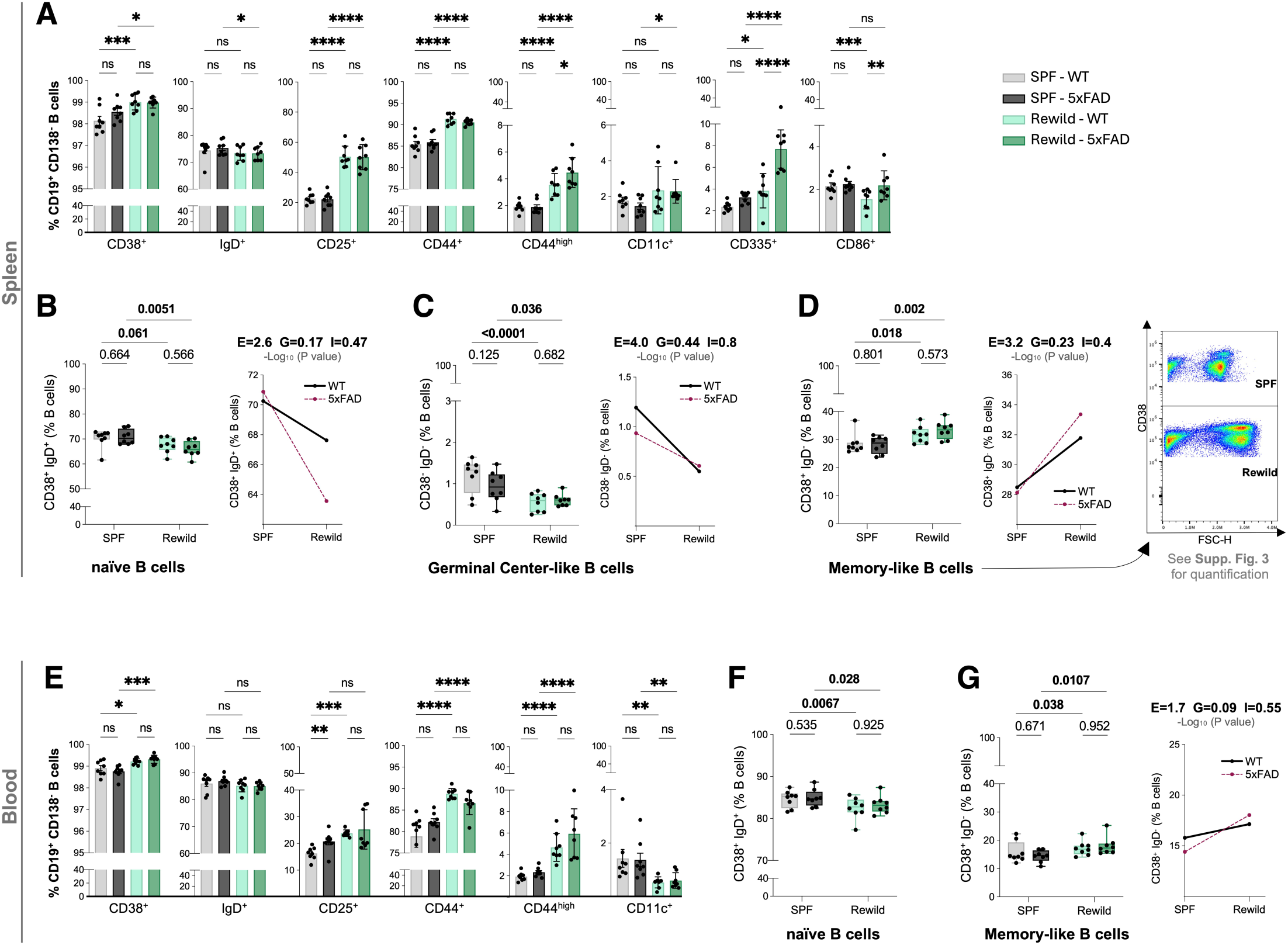
Rewilding increases B cell maturation and activation. Flow cytometry analysis of B-cell phenotypes in spleen (A-D) and blood (E-G) from female SPF-WT, SPF-5xFAD, Rewild-WT, and Rewild-5xFAD mice (N = 8 per group). Gating strategies are provided in Supplementary Figure 1. (A, E) Proportions of marker-positive cells within B lymphocytes are shown for the indicated markers. In spleen (A): CD38, IgD, CD25, CD44, CD44^high^, CD11c, CD335 (NKp46), and CD86. In blood (E): CD38, IgD, CD25, CD44, CD44^high^, and CD11c. Values are expressed as % of total B lymphocytes. (B-D) Splenic B-cell subset proportions expressed as % of B lymphocytes, using an operational CD38/IgD definition: naïve B cells (CD38^+^IgD^+^) (B), germinal center-like B cells (CD38⁻IgD⁻) (C), and memory-like B cells (CD38^+^IgD⁻) (D). The right panel in (D) shows representative CD38 versus FSC plots (SPF vs Rewild) illustrating cell population heterogeneity within the CD38^+^IgD⁻ (memory-like) gate; corresponding quantification of these fractions is provided in Supplementary Figure 3. (F, G) Circulating B-cell subset proportions expressed as % of B lymphocytes: naïve B cells (CD38^+^IgD^+^) (F) and memory-like B cells (CD38^+^IgD⁻) (G). For the boxplot panels (A-G), statistical analyses were performed in R using the same pipeline as described in Methods. The adjacent interaction plots (where shown) were generated independently using a two-way ANOVA full model; E, G, and I denote -log_10_ (*p*) for the main effects of Environment, Genotype, and their Interaction, respectively. \**p* < 0.05, \*\**p* < 0.01, \*\*\**p* < 0.001, \*\*\*\**p* < 0.0001; ns: not significant.

Within the blood compartment, the differentiation shifts were similar to those in the spleen, as rewilding increased the proportion of CD44^+^ and CD44^high^, CD25^+^ and CD38^+^ cells (**Figure 3E**). In addition, circulating B cells downregulated CD11c expression under rewilding (**Figure 3E**). The proportion of IgD⁺ B cells was stable in blood and spleen (**Figure 3E**). Finally, rewilding decreased the proportion of naïve B cells (CD38⁺IgD⁺; **Figure 3F**) and increased that of memory-like B cells (CD38⁺IgD⁻; **Figure 3G**) in blood, consistent with a systemic shift away from naïve phenotypes under rewilding.

### Rewilding expands the ASC compartment and reshapes its differentiation state in spleen and blood

Building on the global expansion of ASCs under rewilding in both spleen and blood of WT and 5xFAD mice (**Figure 2**; **Supplementary Figure 2**), we next asked whether environmental exposure also altered the phenotypic composition of the ASC pool according to CD19 and MHCII markers expression (**Figure 4**). Rewilding significantly increased the number of all ASC subsets in both spleen and blood, independently of genotype (**Figure 4**, count/µL panels).

**Figure 4.**
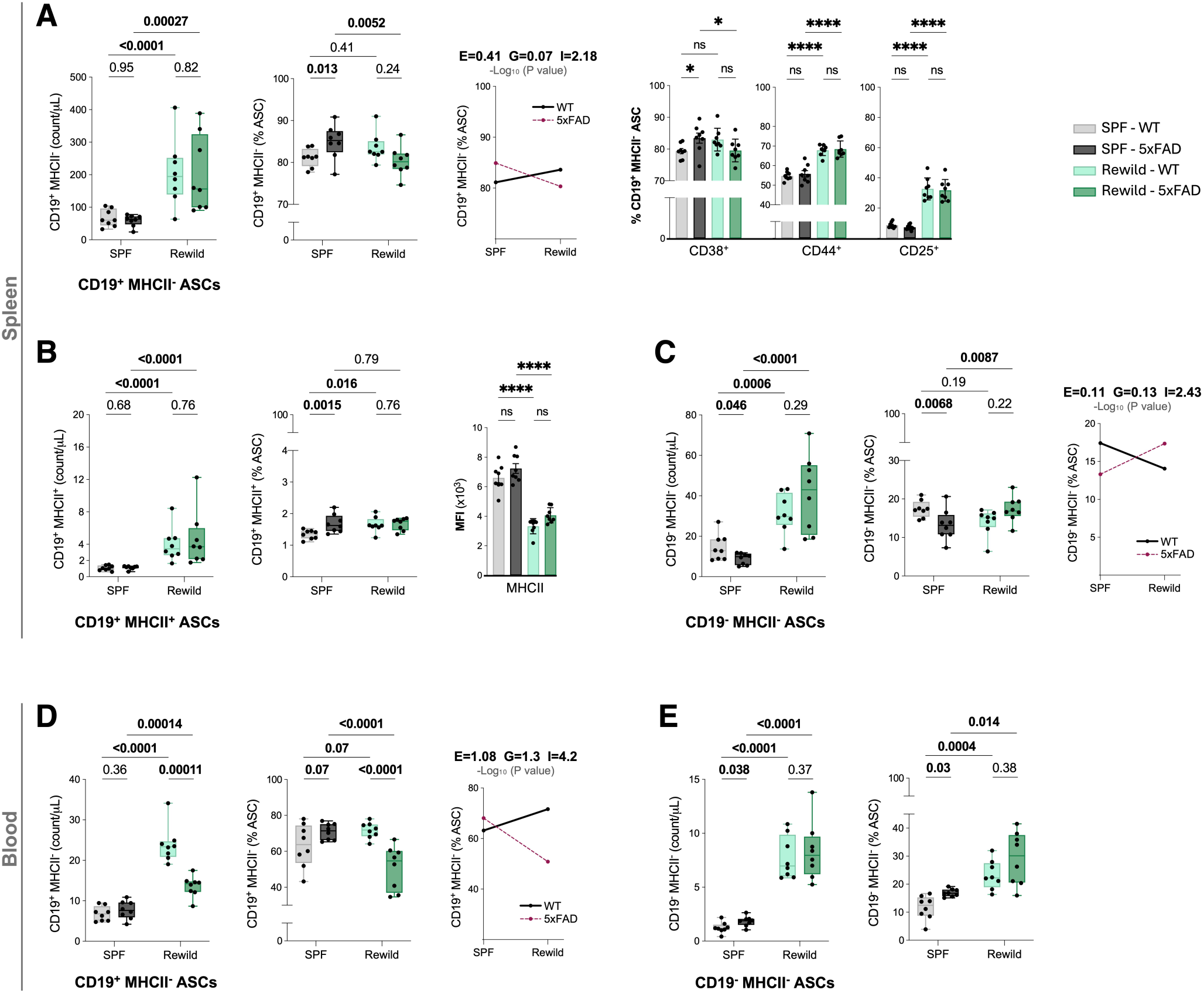
Rewilding expands and promotes differentiation of ASCs. Flow cytometry analysis of splenic (A-C) and circulating ASC subsets (D,E) from female SPF-WT, SPF-5xFAD, Rewild-WT, and Rewild-5xFAD mice (N = 8 per group). Gating strategy of ASC subsets is shown in Supplementary Figure 1. In spleen, the abundant ASC subsets are shown as CD19^+^MHCII^−^ ASCs (A), CD19^+^MHCII^+^ ASCs (B), and CD19^−^MHCII^−^ ASCs (C), whereas in blood the two abundant subsets were CD19^+^MHCII^−^ ASCs (D) and CD19^−^MHCII^−^ ASCs (E). For each subset, data are shown as numbers (counts/µL of acquired sample, absolute abundance) and as proportions of antibody-secreting cells (% ASCs). The interaction plots summarize the full two-way ANOVA model with E, G, and I denoting -log_10_ (*p*) for the main effects of Environment, Genotype, and their Interaction, respectively. Additional phenotypic characterization of splenic CD19^+^MHCII^−^ ASCs is shown in (A) as the proportion of CD38^+^, CD44^+^, and CD25^+^ cells within this subset, and MHCII expression intensity is shown as MFI for splenic CD19^+^MHCII^+^ ASCs in (B). Statistical testing, transformations, and outlier handling were performed using the same pipeline described in Methods. \**p* < 0.05, \*\*\*\**p* < 0.0001; ns: not significant.

Importantly, proportional analyses showed that this expansion was accompanied by a redistribution of ASC states, consistent with enhanced differentiation under rewilding. In the spleen, the proportion of the dominant CD19^+^MHCII^−^ ASCs population decreased selectively in 5xFAD mice, concomitant with a proportional increase in the more differentiated CD19^−^MHCII^−^ subset (**Figure 4A,C**). Thus, in 5xFAD spleen, rewilding was associated not only with expansion of the ASC compartment, but also with a shift toward a more differentiated ASC state. By contrast, the CD19^+^MHCII^+^ splenic subset (plasmablast-like subset) showed only modest proportional changes, but exhibited a marked reduction in MHCII expression intensity under rewilding in both genotypes (**Figure 4B**), consistent with progression away from a less differentiated, antigen-presenting-like ASC phenotype. Further phenotypic characterization of splenic CD19^+^MHCII^−^ ASCs showed that rewilding significantly increased the fraction of CD44^+^ and CD25^+^ cells within this dominant subset (**Figure 4A**), suggesting a more activated or functionally engaged phenotype in both genotypes.

In the blood, this differentiation-associated pattern was even more apparent. Rewilding markedly reduced the proportion of the less differentiated CD19^+^MHCII^−^ subset in 5xFAD mice, alongside a reciprocal enrichment of the more differentiated CD19^−^MHCII^−^ subset (**Figure 4D,E**). Notably, CD19^−^MHCII^−^ ASCs increased both in number and in proportion in both genotypes, compatible with a strong environment-related differentiation effect. To determine whether rewilding-associated remodeling of circulating ASCs translated into altered humoral output, we quantified plasma immunoglobulin isotypes by MSD (**Supplementary Figure 4**). Baseline IgM levels showed a consistent genotype effect (WT > 5xFAD) in both SPF and rewilded conditions, indicating a stable difference in the IgM compartment across environments. In contrast, rewilding selectively shifted isotypes as plasma IgA increased in both genotypes under rewilding, whereas IgG1 decreased under rewilding. IgG2b and IgG2c remained essentially unchanged, consistent with a selective reshaping of humoral output rather than a broad increase across all isotypes (**Supplementary Figure 4**).

Collectively, these results show that rewilding drives not only systemic expansion and differentiation remodeling of the ASC compartment, but also a selective reshaping of circulating humoral output, with particularly evident genotype-dependent redistribution of ASC states in 5xFAD mice.

### Rewilding reshapes peripheral T-cell differentiation states

Because CD62L was not included in the T-cell panel, CD8⁺ T-cell differentiation states were defined operationally using a hierarchical gating strategy (**Supplementary Figure 1**). After lineage exclusion and selection of CD8a⁺ T cells, KLRG1 was used to separate KLRG1⁺ short-lived effector-like CD8⁺ cells (SLECs) from the KLRG1⁻ compartment. Within KLRG1⁻ CD8⁺ cells, we further focused on a CD127⁺CD27⁺ precursor-enriched pool and partitioned this compartment by CD44 intensity using an FMO-based CD44 threshold to define CD44⁻ (naïve), CD44⁺ (virtual memory), and CD44^high^ (memory precursor effector cells, MPECs) fractions. This CD44-based stratification captured a prominent environment-associated shift as rewilded mice displayed a clear rightward displacement of the CD44 distribution, with fewer CD44⁻ events and enrichment of CD44⁺, compared to SPF controls (**Figure 5A**).

**Figure 5.**
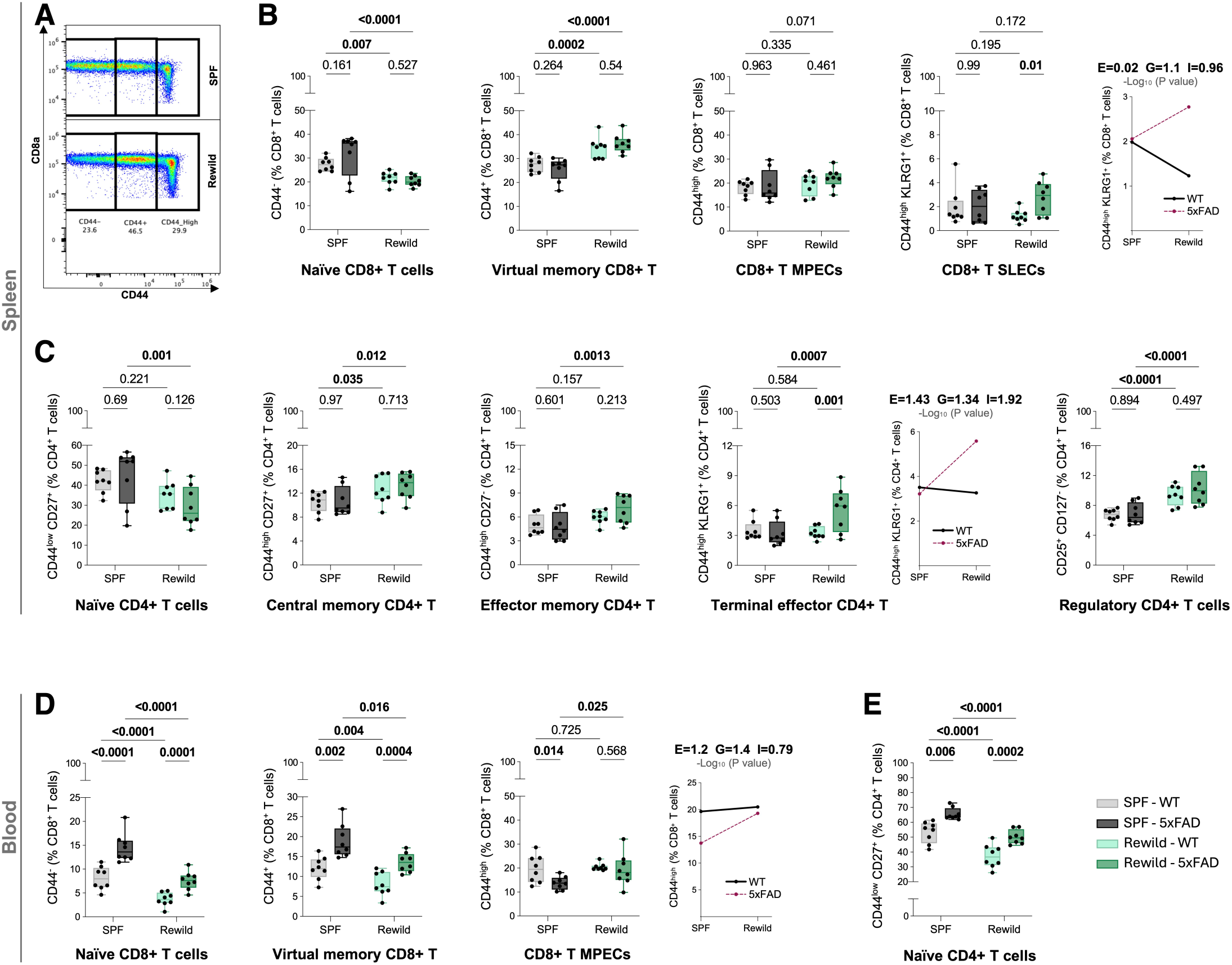
Rewilding shifts circulating and splenic T-cell differentiation states. Flow cytometry analysis of T-cell subsets in spleen (A-C) and blood (D-E) from female SPF-WT, SPF-5xFAD, Rewild-WT, and Rewild-5xFAD mice (N = 8 per group). Gating strategies for subset definitions are provided in Supplementary Figure 1. (A) Representative CD8a versus CD44 plots illustrating the gating of splenic CD8⁺ T-cell subsets and highlighting a global shift toward higher CD44 expression in rewilded mice relative to SPF controls. (B) Splenic CD8⁺ T-cell subset proportions (% of CD8^+^ T cells): naïve CD8⁺ T-cell, virtual memory CD8⁺ T-cell, memory precursor effector cells (CD8⁺ MPECs), and short-lived effector cells (CD8⁺ SLECs). (C) Splenic CD4⁺ T-cell subset proportions (% of CD4^+^ T cells): naïve CD4⁺ T-cell, central memory CD4⁺ T-cell, effector memory CD4⁺ T-cell, terminal effector CD4⁺ T-cell, and regulatory CD4⁺ T-cells. (D) Blood CD8⁺ T-cell subset proportions (% of CD8^+^ T-cells): naïve CD8⁺ T-cell, virtual memory CD8⁺ T-cell, and CD8⁺ MPECs. (E) Blood naïve CD4⁺ T cells shown as % of CD4^+^ T-cells (other CD4⁺ differentiation subsets were too sparse for robust quantification in blood). Interaction plots (where shown) and statistical testing were performed as described in Methods.

In the spleen, rewilding shifted the CD8⁺ compartment away from naïve states and toward antigen-experienced phenotypes (**Figure 5B**). The naïve CD8⁺ proportion decreased in both genotypes, while the virtual memory proportion increased, consistent with the CD44 expression shift observed in representative plots. The intermediate precursor (MPECs) and most differentiated CD8⁺ effector subset (SLECs) showed modest changes in proportion across environments, indicating that rewilding impacts early CD8⁺ differentiation balance within secondary lymphoid tissue (**Figure 5B**). CD4⁺ differentiation in spleen exhibited a genotype-selective remodeling pattern (**Figure 5C**). Rewilding reduced naïve CD4⁺ representation while increasing central memory and effector-associated CD4⁺ proportion in 5xFAD mice, consistent with an overall state more prone to shift toward more antigen-experienced CD4⁺ states in AD mice. In contrast, regulatory CD4⁺ T cells (CD25⁺CD127⁻) increased strongly in spleen under rewilding (**Figure 5C**) in both genotypes, indicating that rewilding robustly expands an immunoregulatory population.

In the blood, the rewilding-driven shift was particularly pronounced for CD8⁺ T cells (**Figure 5D**). Rewilding strongly depleted the naïve CD8⁺ and the virtual memory fractions in both genotypes, along with a modest increase in MPECs in 5xFAD mice. Notably, genotype differences, apparent under SPF conditions, were compressed under rewilding, as both genotypes shifted toward more antigen-experienced CD8⁺ cells (**Figure 5D**). In the circulating CD4⁺ T cells compartment, the principal detectable differentiation readout, naïve CD4⁺, was substantially reduced under rewilding (**Figure 5E**), mirroring the broader systemic shift away from naïve phenotypes. Together, these data show that indoor rewilding drives a consistent, tissue-dependent shift in T-cell differentiation from naïve toward antigen-experienced states.

### Rewilding induces coordinated phenotype switching across peripheral innate myeloid compartments

To assess whether rewilding reprograms innate immune states, we quantified major splenic and circulating myeloid compartments and tracked their activation-marker profiles. A prominent feature of the rewilded spleen was the expansion of a distinct SSC^low^ neutrophil state (switched neutrophils) within the Ly6G⁺ compartment (**Figure 6A**). This fraction increased strongly under rewilding (p<0.0001), whereas the conventional SSC^high^ neutrophil compartment showed the reciprocal shift (**Figure 6B**; p<0.0001). Importantly, this reconfiguration was accompanied by a coordinated marker remodeling consistent across both neutrophil states as indicated by pronounced environment-associated shifts in CD86, CD44, CD38, and CD25 MFIs. Markedly, SSC^low^ neutrophils were bright in CD38 and CD25 expression relative to SSC^high^ neutrophils, and showed decreased CD44 expression in rewilding conditions compared to SSC^high^ neutrophils (**Figure 6A, B**).

**Figure 6.**
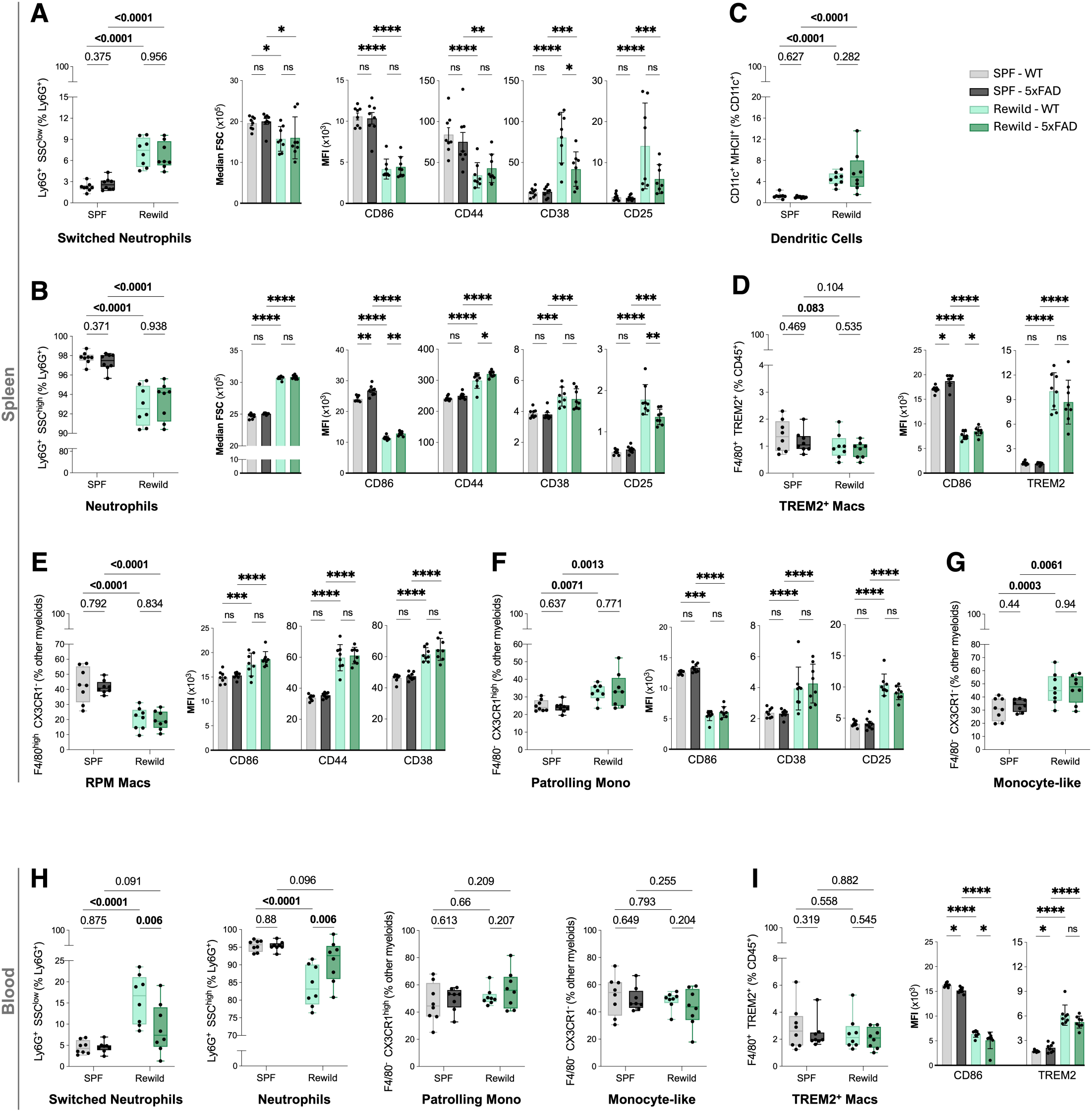
Rewilding remodels myeloid phenotypes and activation-marker expression in spleen and blood. Flow cytometry analysis of myeloid subsets in spleen (A-G) and blood (H-I) from female SPF-WT, SPF-5xFAD, Rewild-WT, and Rewild-5xFAD mice (N = 8 per group). Gating strategies are provided in Supplementary Figure 1. (A) Switched neutrophils in spleen, quantified as Ly6G⁺SSC^low^ (% of Ly6G⁺), and (B) Splenic neutrophils quantified as Ly6G⁺SSC^high^ (% of Ly6G⁺), along with their median-FSC and marker expression (MFI) for CD86, CD44, CD38, and CD25. (C) Splenic dendritic cells (DCs) quantified as CD11c⁺MHCII⁺ (% of CD11c⁺). (D) Splenic TREM2⁺ macrophages quantified as F4/80⁺TREM2⁺ (% of CD45⁺), with marker expression (MFI) for CD86 and TREM2. (E) Splenic red pulp macrophages (RPM Macs) quantified as F4/80^high^CX3CR1⁻ (% of other myeloids), with marker expression (MFI) for CD86, CD44, and CD38. (F) Splenic patrolling monocytes quantified as F4/80⁻CX3CR1^high^ (% of other myeloids), with marker expression (MFI) for CD86, CD38, and CD25. (G) Splenic monocyte-like cells quantified as F4/80⁻CX3CR1⁻ (% of other myeloids). (H) Blood myeloid subsets: switched neutrophils, neutrophils, patrolling monocytes and monocyte-like cells. (I) Blood TREM2⁺ macrophages quantified as F4/80⁺TREM2⁺ (% of CD45⁺), with marker expression (MFI) for CD86 and TREM2. Statistical testing, transformations, and outlier handling were performed as described in Methods. \**p* < 0.05, \*\**p* < 0.01, \*\*\**p* < 0.001, \*\*\*\**p* < 0.0001; ns: not significant.

Beyond neutrophils, rewilding drove broad innate activation marker remodeling across splenic antigen-presenting cells and monocytes/macrophages. Dendritic cells (DCs; CD11c⁺MHCII⁺) increased under rewilding (**Figure 6C**). Splenic red pulp macrophages (RPM; F4/80^high^CX3CR1⁻) strongly decreased in proportion (**Figure 6E**) and increased in CD86/CD44/CD38 expression, consistent with major state remodeling within this macrophage niche (**Figure 6E**). Similarly, splenic patrolling monocytes increased (F4/80⁻CX3CR1^high^; **Figure 6F**) and showed strong rewilding-associated increases in CD38/CD25 expression while CD86 expression strongly decreased, consistent with an activation-marker remodeling. Further, the broader monocyte-enriched compartment also increased in rewilding conditions (**Figure 6G**).

A notable pattern emerged for TREM2⁺ macrophages (F4/80^+^SSC^high^TREM2^+^): while their proportion among leukocytes did not change across environments (**Figure 6D**), their phenotype shifted substantially. CD86 expression decreased while TREM2 expression increased under rewilding (**Figure 6D**), indicating a robust marker-defined reprogramming of this cell population.

In blood, the same phenotypic switch was detectable but was generally more modest in magnitude. The SSC^low^ switched neutrophils increased in proportion under rewilding while the SSC^high^ conventional neutrophils decreased (**Figure 6H**). Other circulating myeloid cells showed weaker or non-significant environment effects on frequency (**Figure 6H**). Notably, circulating TREM2⁺ macrophages again showed strong phenotype remodeling with substantially decreased CD86 and increased TREM2 expression despite limited changes in overall number (**Figure 6I**).

Taken together, peripheral immunophenotyping indicates that indoor rewilding broadly reshapes both adaptive and innate immune compartments in spleen and blood, with expansion of B-lineage and ASC populations, shifts in T-cell differentiation, and coordinated phenotypic changes across myeloid lineages.

### Rewilding remodels brain microglial state balance and reprograms gene expression

These consistent systemic effects in both WT and 5xFAD mice motivated us to next examine whether analogous remodeling was detectable within the brain. Using P2RY12 to stratify brain microglia (**Figure 7A**), we quantified marker profiles in CX3CR1^high^P2RY12^high^ (homeostatic-like) versus CX3CR1^high^P2RY12^dim^ (reactive-like) microglial states (**Figure 7B, C**). Within both states, rewilding was associated with a strong decrease in CX3CR1 and CD86 expression, and an increase in CD38 and CD25 MFIs, in both genotypes (**Figure 7B, C**). To minimize sensitivity to dissociation-related variation in total immune-cell yield, we assessed a homeostatic-like/reactive-like ratio within the CX3CR1^high^ compartment (**Figure 7D**). Rewilding increased this ratio within WT, indicating a more homeostatic fraction relative to SPF-WT, consistent with WT-specific increase in P2RY12 MFI in rewilding conditions (**Figure 7B, C**).

**Figure 7.**
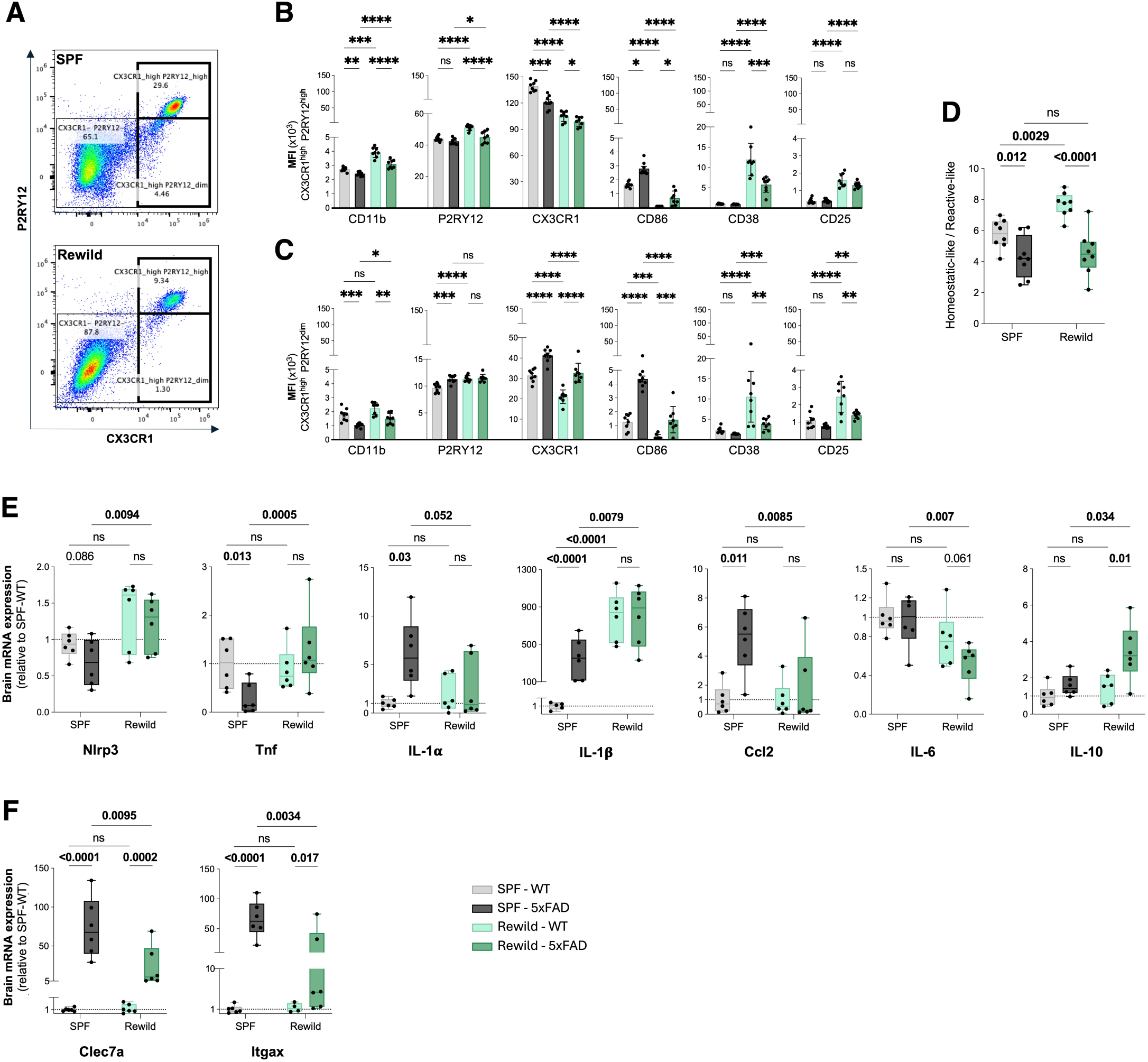
Rewilding remodels brain microglial state balance and reprograms gene expression. Flow cytometry and qRT-PCR analyses were performed on brain tissue from female SPF-WT, SPF-5xFAD, Rewild-WT, and Rewild-5xFAD mice (N = 8 per group). The brain flow cytometry gating strategy is shown in Supplementary Figure 5. (A) Representative bivariate plots (P2RY12 vs CX3CR1) illustrating the microglial gating approach under SPF and Rewild conditions and the distribution of CX3CR1/P2RY12-defined subsets. (B) Microglial marker expression (MFI) for CD11b, P2RY12, CX3CR1, CD86, CD38, and CD25 in CX3CR1^high^P2RY12^high^ cells (homeostatic-like microglia). (C) The same markers in CX3CR1^high^P2RY12^dim^ cells (reactive-like microglia). (D) Homeostatic-like/Reactive-like ratio metric was used to minimize sensitivity to between-sample variation in total brain immune cell yield introduced during tissue dissociation and processing. (E-F) Brain mRNA expression measured by qRT-PCR for *Nlrp3*, *Tnf*, *IL-1α*, *IL-1β*, *Ccl2*, *IL-6*, and *IL-10* (E), and for *Clec7a*, *Itgax* (F), normalized to housekeeping gene *Gapdh*. Data are displayed as fold change relative to the SPF-WT group. Statistical testing was performed on log_2_-transformed fold-change values using a two-way ANOVA model with Šidák’s correction. MFI values were analyzed as described in Methods. \**p* < 0.05, \*\**p* < 0.01, \*\*\**p* < 0.001, \*\*\*\**p* < 0.0001; ns: not significant.

In addition, qRT-PCR revealed robust rewilding-associated remodeling of inflammatory gene expression in 5xFAD mice (**Figure 7E**). With the exception of *IL-1β*, where the rewilding effect was significant in both genotypes, several other transcripts showed a strong environmental effect specific to 5xFAD mice. Indeed, in contrast to SPF-5xFAD mice, Rewild-5xFAD mice showed a distinct inflammatory profile, marked by reduced *IL-1α*, *Ccl2*, and *IL-6* transcripts alongside increased *IL-10*, *Nlrp3*, and *Tnf* expression (**Figure 7E**), suggesting a transition from a classical neuroinflammatory state toward a more surveillant, homeostatic-like, yet potentially primed microglial phenotype. Similarly, two microglial disease-associated markers, Clec7a and Itgax, were strongly reduced in 5xFAD mice under rewilding conditions (**Figure 7F**), consistent with an attenuation of this DAM-related transcriptional axis.

### Rewilding modulates microglia-Aβ interaction and metabolism

We next examined whether the microglial state shifts observed by flow cytometry and qRT-PCR were reflected in microglial morphology and in Aβ plaque-associated microglial physical interactions *in situ*. Iba1^+^ microglia were classified into three morphological states (homeostatic, reactive, and amoeboid) based on soma size and ramification complexity, as illustrated by representative confocal images (**Figure 8A**). Quantitative classification revealed region-specific effects of genotype and environment on microglial state distribution (**Figure 8B**).

**Figure 8.**
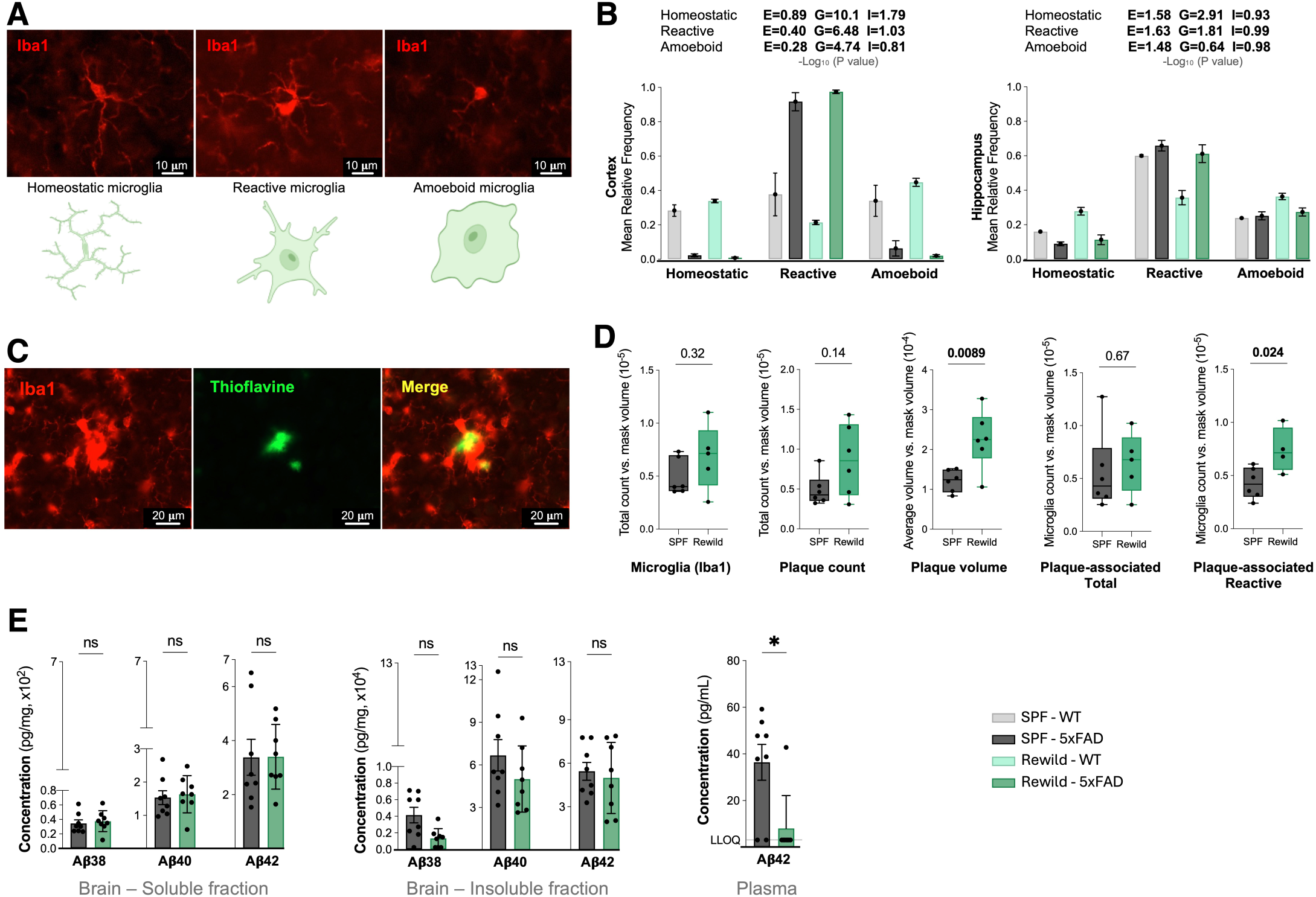
Rewilding modulates microglia-Aβ interaction and metabolism. Histological and biochemical analyses were performed in female SPF-WT, SPF-5xFAD, Rewild-WT, and Rewild-5xFAD mice (N = 6–8 per group, as indicated). Microglia were visualized by Iba1 immunostaining and amyloid plaques by Thioflavin S staining. (A) Representative Iba1 images illustrating microglial morphological states used for classification: Homeostatic, Reactive, and Amoeboid. Scale bar, 10 µm. (B) Quantification of microglial morphological states shown as mean cell frequency in cortex and hippocampus (right hemisphere fixed; N = 6 per group). Insets report -log_10_ (*p*) for the main effects of Environment (E) and Genotype (G) and their Interaction (I) from the full two-factor ANOVA model, as indicated in Methods. (C) Representative images showing microglia and amyloid plaques with Iba1 (microglia), Thioflavin S (plaques), and merged channels. Scale bar, 20µm. (D) Quantification of total microglial signal (Iba1), plaque count, plaque volume, and plaque-associated microglial measures, total and reactive, as indicated. (right hemisphere fixed; N = 6 per group). (E) Human Aβ peptide quantification by MSD immunoassays in brain and plasma (N = 8 per group). Brain concentrations are expressed as pg/mg of total proteins; plasma Aβ_42_ is reported in pg/mL. Statistical testing was performed on log_10_-transformed data. Values below the lower limit of quantitation (LLOQ) are indicated where applicable. Plasma Aβ_40_ was bellow detection range for all samples (not shown). Two-group comparisons used two-tailed t-tests or Mann-Whitney tests as appropriate. Exact *p*-values and significance annotations are shown on the plots (\**p* < 0.05; ns, not significant).

In the cortex, morphology was primarily shaped by genotype. Relative to WT mice, 5xFAD animals showed a marked loss of homeostatic microglia together with a strong enrichment in reactive microglia, consistent with amyloid-associated activation (**Figure 8B**). Cortical homeostatic microglia were significantly reduced in both SPF-5xFAD and Rewild-5xFAD mice compared with both WT groups (all *p* < 0.0001), while reactive microglia were significantly increased in both 5xFAD groups relative to both WT groups (all *p* < 0.0001). Amoeboid microglia were also lower in 5xFAD mice than in WT mice (all *p* < 0.001). Rewilding had only a limited effect in the cortex, with a modest increase in the homeostatic fraction in WT mice (*p*=0.0241), but no significant effect within 5xFAD animals. In the hippocampus, microglial morphological states were influenced by both environment and genotype depending on the phenotype considered, whereas no significant G × E interaction was detected for any morphological class (**Figure 8B**). Thus, although group mean differences were visually apparent, these patterns could not be formally attributed to genotype-specific responses to the environmental condition. We therefore focused subsequent analyses on the cortex, where microglial remodeling was most robust.

Dual-labeling for Iba1 and Thioflavin S using high-resolution confocal imaging revealed a close spatial association between microglia and Aβ plaques, suggesting increased phagocytosis (**Figure 8C**). Quantitative 3D analysis of confocal z-stacks revealed that total microglial numbers per tissue volume were comparable between SPF-5xFAD and Rewild-5xFAD mice (**Figure 8D**). However, while total Aβ plaque counts remained unchanged, rewilding significantly increased their mean volume (**Figure 8D**), suggesting altered plaque dynamics or remodeling. Importantly, the density of reactive, plaque-associated microglia was significantly higher in 5xFAD-Rewild mice (**Figure 8D**). MSD quantification of human Aβ peptides showed that brain soluble and formic-acid insoluble Aβ_38/40/42_ levels were broadly similar across groups, whereas plasma Aβ_42_ was significantly reduced under rewilding (**Figure 8E**).

These findings suggest that rewilding alters microglial engagement and morphology in ways that support altered metabolic and surveillance functions. Microglia in naturalized environments seem to maintain a balanced state between resting and reactive modes, possibly enabling continuous plaque remodeling without worsening neuroinflammation. Overall, these results demonstrate how environmental microbial diversity could influence microglial structure and Aβ interactions, supporting a model where rewilding encourages adaptive immune-metabolic regulation rather than outright activation.

### Rewilding redirects brain transcriptional signatures toward human AD

To investigate how microbial exposure reshapes the brain transcriptome, we performed whole-brain Nanopore sequencing in SPF-WT, SPF-5xFAD, Rewild-WT, and Rewild-5xFAD mice, along with wild *Mus musculus* brains used as a natural reference (**Supplementary File 1**). Strikingly, indoor rewilding induced extensive transcriptional remodeling, with a 61% overlap of significant (± 1.25-fold-change) differentially expressed genes (DEGs) between Rewild-WT and Wild-caught (Feral) mice (odds ratio = 2.94, *p*=4.9×10⁻^219^) when compared to SPF mice, demonstrating strong convergence of CNS gene expression toward wild-relevant physiological states (**Figure 9A**). Correlation analyses across differentially expressed gene contrasts further revealed that rewilding substantially redirected brain transcriptional signatures, driving convergence across biological contexts. Rewild-WT mice shared a modest but significant correlation with Wild (Feral) mice (Pearson r=0.363, *p*=1.04×10⁻^62^; n=1984 genes), indicating that environmental enrichment alone is sufficient to partially recapitulate feral transcriptional signatures (**Figure 9B**). Strikingly, the transcriptional profile of Rewild-WT mice showed a stronger correlation with SPF-5xFAD mice (r=0.748, *p*=1.03×10⁻^84^; n=467), suggesting that rewilding and Alzheimer’s-related pathology engage overlapping gene expression signatures (**Figure 9C**). This convergence was most pronounced in Rewild-5xFAD mice, which exhibited a near-complete correlation with Rewild-WT mice (r=0.962, *p* <1×10⁻^300^; n=1976), demonstrating that the rewilding transcriptional signature dominates and is highly consistent regardless of genotype (**Figure 9D**).

**Figure 9.**
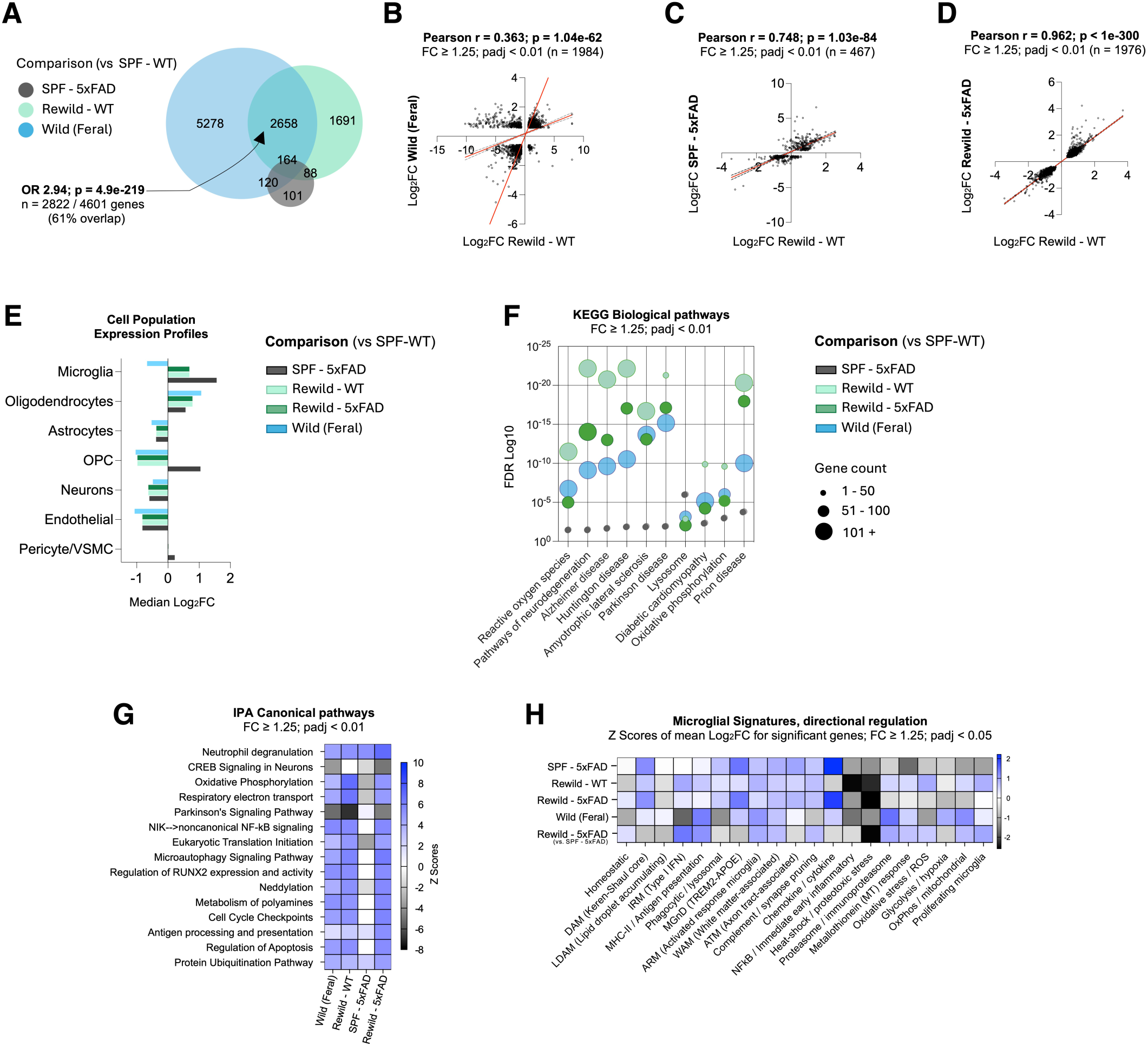
Rewilding substantially redirects brain transcriptional signatures. Nanopore sequencing was performed on whole left brain hemispheres from female mice (N = 6 per group): SPF-WT, SPF-5xFAD, Rewild-WT, Rewild-5xFAD, and Wild (Feral). Differential expression was assessed using DESeq2. Unless indicated, significance thresholds are FC ≥ 1.25 (|log_2_FC| ≥ ∼0.32) and *p*adj < 0.01. (A) Venn diagram showing overlap of significantly regulated genes across comparisons vs SPF-WT for SPF-5xFAD, Rewild-WT, and Wild (Feral). The number of significant genes in each comparison and overlap sizes are shown. Statistics for the largest overlap are reported. (B–D) Pearson correlation of log_2_FC values among shared significant genes (FC ≥ 1.25; *p*adj < 0.01), comparing Rewild-WT with Wild (Feral) (B), SPF-5xFAD (C), and Rewild-5xFAD (D). Sample sizes (n), Pearson r, and *p*-values are indicated in each panel. (E) Cell population expression profiles summarizing the median log_2_FC of significant genes across comparisons versus SPF-WT for major brain cell populations (Microglia, Oligodendrocytes, Astrocytes, OPC, Neurons, Endothelial, Pericyte/Vascular Smooth Muscle Cells). Bar shading indicates comparison group. (F) Bubble plot of enriched KEGG biological pathways across comparisons (FC ≥ 1.25; *p*adj < 0.01). Bubble color indicates the comparison group, and the y-axis shows enrichment significance (FDR log_10_). Dot size encodes the number of significant genes (1–50, 51–100, or ≥101) contributing to each pathway. (G) Heatmap of IPA canonical pathways across comparisons (FC ≥ 1.25; *p*adj < 0.01), displayed as IPA activation Z-scores. (H) Directional regulation of curated microglial gene signatures across mouse contrasts, shown as Z-scores of mean log_2_FC for significant genes, showing coordinated shifts in DAM, phagocytic/lysosomal, antigen-presentation, and metabolic programs in rewilded 5xFAD mice.

We next quantified transcriptional changes of canonical markers (genes) across all major CNS lineages^46–51^ (**Supplementary File 2**), where both environment and genotype exerted distinct effects relative to SPF-WT mice **(Figure 9E)**. SPF-5xFAD mice exhibited pronounced microgliosis, whereas rewilding significantly attenuated microglial activation. Oligodendrocytes, astrocytes, and endothelial cells showed moderate, directionally consistent remodeling, although OPCs were uniquely elevated in SPF-5xFAD. Pericytes and vascular smooth muscle cells (VSMC) remained largely unchanged across groups.

Pathway enrichment using KEGG (**Figure 9F**) and Ingenuity Pathway Analysis (IPA) (**Figure 9G**) revealed strong induction of immune, mitochondrial, oxidative phosphorylation, and antigen-presentation pathways in rewilded mice (**Supplementary File 3**). Notably, Rewild-5xFAD displayed substantially higher pathway activation scores than SPF-5xFAD, indicating synergistic effects of environmental complexity and amyloid pathology. Interestingly, among the top-scoring IPA “upstream regulators” of these pathways are immune (e.g., immunoglobulins) and AD (e.g., MAPT, APP, and PSEN1)-related genes (**Supplementary File 3**).

Quantitative gene expression analysis of canonical and emerging microglial states^52–54^ reveals that SPF 5xFAD mice strongly activate gene signatures associated with DAM, as expected, alongside signatures linked to other disease-related microglial states, including microglia with a neurodegenerative phenotype (MGnD), axon tract-associated microglia (ATM), and white matter-associated microglia (WAM), as well as phagocytic and lysosomal programs, while concomitantly losing homeostatic microglial signatures (**Figure 9H**). In contrast, rewilded WT (and Wild) mice also engage DAM-like programs but display stronger induction of antigen presentation, proteasomal, oxidative stress, and metabolic microglial states, consistent with immune conditioning. Importantly, in rewilded 5xFAD mice, the degenerative microglial landscape was more diversified and attenuated, rather than reflecting canonical DAM amplification.

The expression analyses of individual AD-associated genes revealed that rewilding regulated several key human risk loci (e.g., *Apoe*, *Trem2*, *Clu*, and *Picalm*) (**Figure 10A-E**), strengthening the connection between ecological exposure and disease-relevant transcriptional signatures. The modulatory effects of rewilding on selected microglial (*Iba1*) and astrocytic (*Gfap*) core genes could be reproduced at both mRNA and protein levels (**Figure 10B, D**).

**Figure 10.**
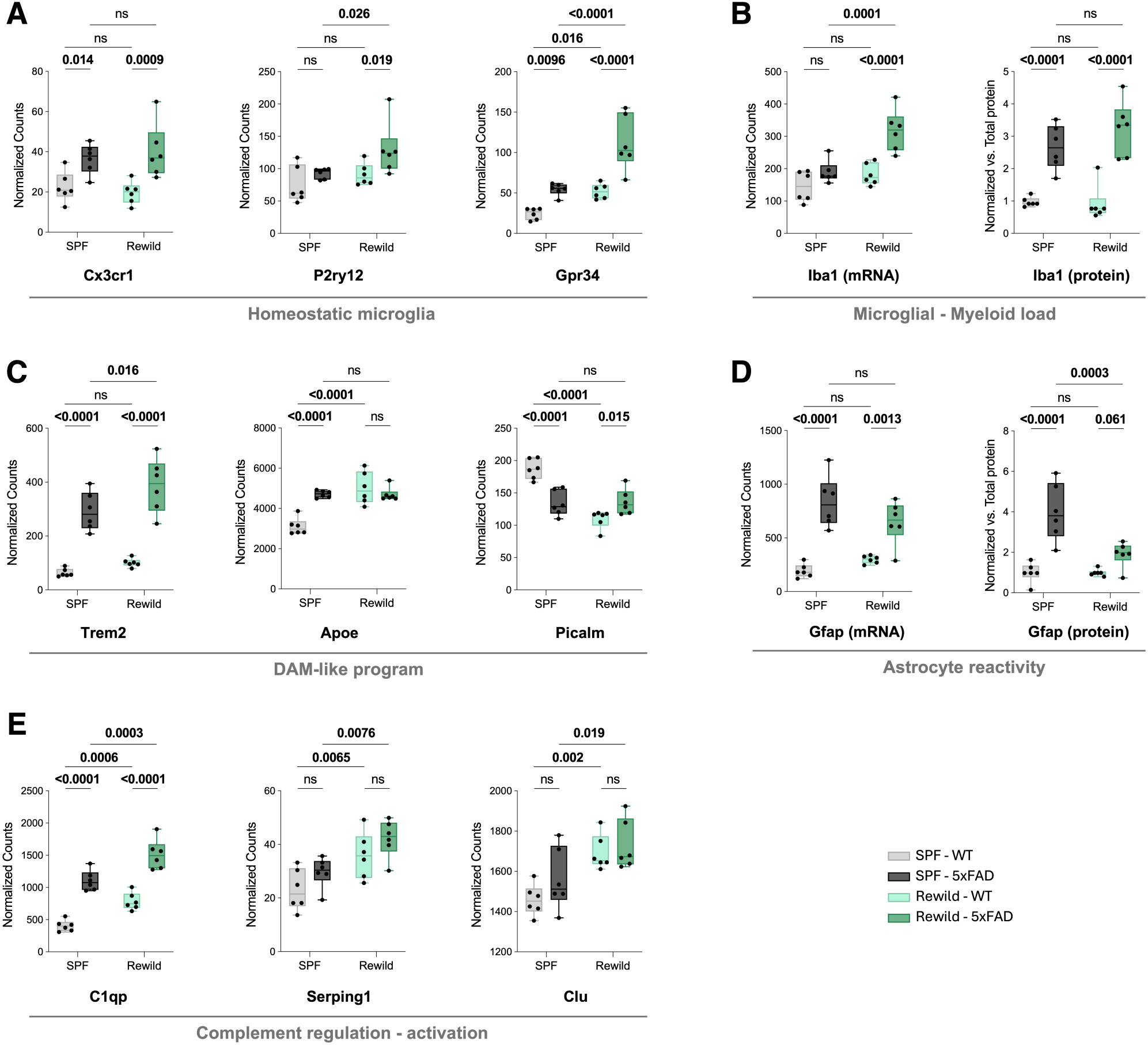
Rewilding modulates microglial and AD-associated gene signatures in the brain. Nanopore sequencing and immunoblot analyses were performed on brain tissue from female SPF-WT, SPF-5xFAD, Rewild-WT, and Rewild-5xFAD mice (N = 6 per group). For Nanopore-seq panels, expression is shown as DESeq2-normalized counts. For immunoblots, ECL signal intensities were normalized to total protein (Ponceau staining). (A) Homeostatic microglial gene expression: *Cx3cr1*, *P2ry12*, and *Gpr34*. (B) Microglial/myeloid load assessed by Nanopore-seq *Aif1* (Iba1) expression and Iba1 protein levels (immunoblot). (C) Microglial activation/DAM-like genes: *Trem2*, *Apoe*, and *Picalm*. (D) Astrocyte reactivity assessed by Nanopore-seq *Gfap* expression and Gfap protein levels (immunoblot). (E) Complement-related genes: *C1qp*, *Serping1*, and *Clu*. For Nanopore-seq panels, *p*-values were derived from DESeq2 model contrasts. For immunoblot quantification, group comparisons were performed using a two-way repeated measures ANOVA with Šidák’s multiple comparison test.

Notably, cross-species pathway integration using IPA showed that rewilded 5xFAD mice, but not their SPF counterparts, exhibited transcriptional profiles that more closely aligned with bulk RNA-Seq datasets from human AD brain^55^, particularly in the entorhinal cortex and hippocampus (**Figure 11D**). This effect was not observed in the postcentral gyrus or superior frontal gyrus from AD individuals (**Supplementary File 3**), suggesting region-specific and Aβ-prone effects. Importantly, this increased human concordance was paralleled by engagement of the human Alzheimer microglia (HAM) signature^56^, a human AD-enriched microglial transcriptional program that captures disease-relevant microglial features not fully encompassed by canonical murine DAM transcriptional program. Largely absent under SPF conditions, this HAM signature was strongly induced in rewilded and wild mice, together with increased overlap with bulk human AD brain expression profiles (**Figure 11E**). Collectively, these findings suggest that environmental microbial complexity reorients the murine CNS toward a transcriptional state, and specifically a microglial response program, that more closely resembles human AD.

**Figure 11.**
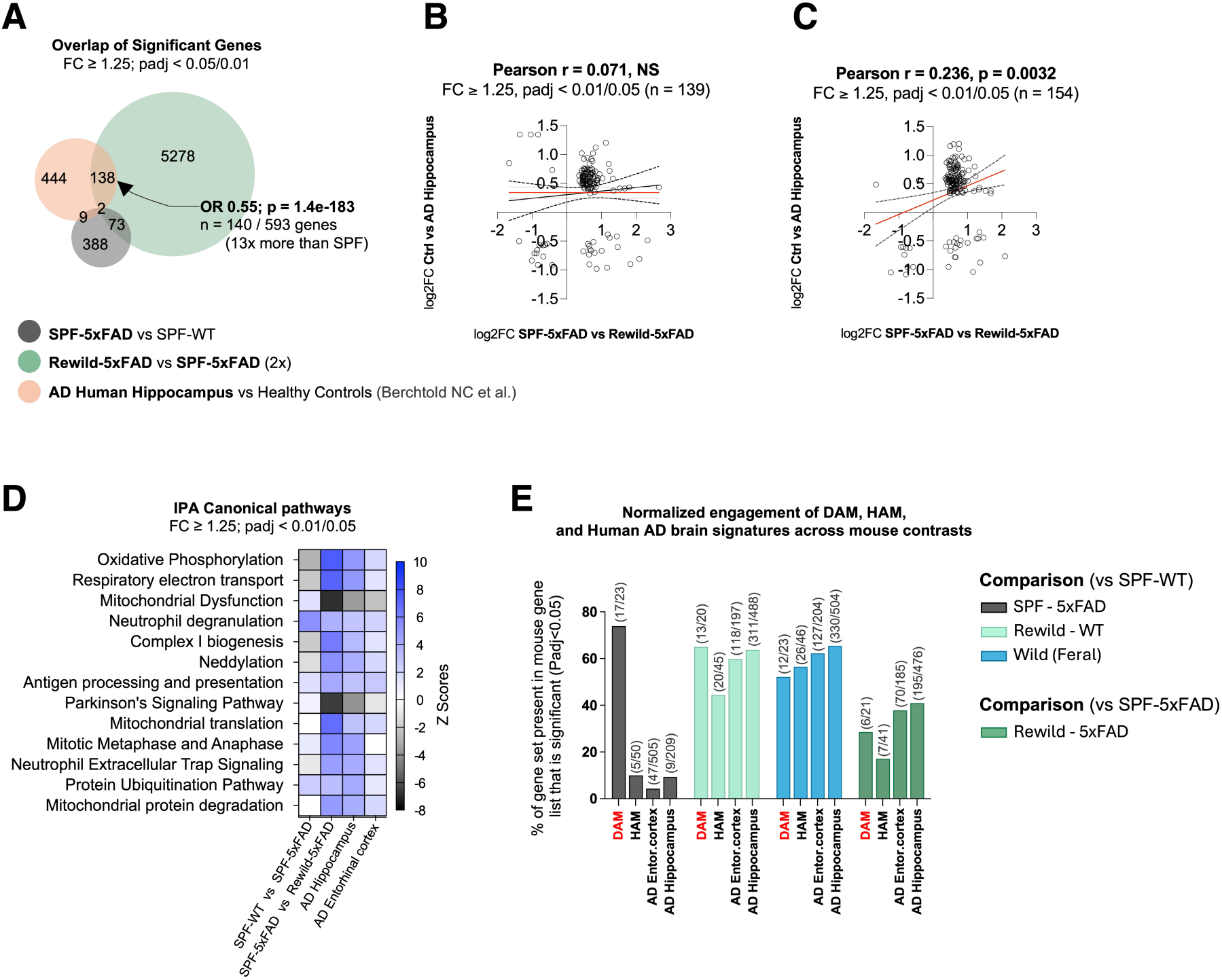
Rewilding enhances transcriptional convergence between 5xFAD mouse brains and human AD. (A) Overlap of differentially expressed genes (DEGs) between Rewild-5xFAD vs SPF-5xFAD (two-fold contrast) and human AD hippocampus (Berchtold Nat Commun). Rewilded 5xFAD mice show a markedly larger overlap (140 shared DEGs; OR = 0.55, *p* = 1.4 × 10⁻^183^) representing a 13-fold increase compared with the overlap observed under SPF conditions (SPF-5xFAD vs SPF-WT). (B) Cross-species comparison of log_2_ fold changes between SPF-5xFAD and human AD hippocampus, showing no meaningful correlation with human AD hippocampus (Pearson r = 0.071, NS; n = 139 DEGs), highlighting the weak translational relevance of conventional SPF housing. (C) A significant correlation is observed (Pearson r = 0.236, *p* = 0.0032; n = 154 DEGs) between rewilded 5xFAD mice and the human AD hippocampus. (D) Canonical pathway analysis (IPA) of DEGs from SPF-5xFAD, Rewild-5xFAD (2x), and two human AD regions (hippocampus and entorhinal cortex). Rewilding amplifies the enrichment of pathways strongly implicated in human AD, including Oxidative Phosphorylation, Respiratory Electron Transport, Mitochondrial Dysfunction, Complex I biogenesis, Protein Ubiquitination, Neutrophil Degranulation/NETs, and Antigen Processing and Presentation, bringing the mouse transcriptional signature closer to human disease. Z-scores reflect pathway directionality across species. (E) Normalized engagement of DAM, HAM, and human AD brain transcriptional signatures across mouse contrasts, expressed as the percentage of signature genes significantly regulated, showing increased overlap with human AD signatures in rewilded mice.

## DISCUSSION

This study demonstrates that ecological exposure profoundly restructures both peripheral and brain immune landscapes in WT and 5xFAD mice. By bridging the immunological sterility of SPF housing with lifelike microbial and antigenic complexity, our indoor rewilding approach restores key components of immune maturation, spanning effector, regulatory, and innate networks, and redefines the neuroimmune tone underlying Aβ-driven pathology. Together, these results establish dirty mouse modeling as an alternative, robust experimental framework for investigating how environmental complexity modulates systemic-neuroimmune crosstalk and disease progression, and for improving the translational fidelity of preclinical neurodegenerative disease research.

Our indoor rewilding platform^37^, which bridges the concepts of (outdoor) rewilding and naturalization^12,44,57,58^, offers a controlled ecosystem to promote natural microbial exposure in laboratory mice. In line with previous dirty mouse models^5,19^, our indoor rewilding recalibrated peripheral B- and T-cell populations toward phenotypes characteristic of immunological maturity, including expanded transitional and activated B cells, increased antibody-secreting cells, an enriched splenic CD4⁺ Treg subset, and broad phenotypic shifts across differentiation and activation markers. This environmental “training” effectively corrects the immune immaturity typical of sterile SPF housing, moving the host from a naïve state toward antigen-experienced readiness^6,9,19,20^. Notably, the dominant CD8⁺ T-cell profile observed in SPF-5xFAD mice was normalized to WT levels under rewilding, accompanied by an enhanced CD4⁺/CD8⁺ T-cell ratio in the blood and expansion of splenic Tregs. This suggests that a complex environment acts as a biological buffer, attenuating the extreme immune skewing driven by amyloid-related genetic mutations under isolated laboratory conditions, as shown in other genetic mouse models^17^.

This adaptive immune recalibration was paralleled by a coordinated state-switch across innate immune cell populations, shifting them toward surveillance and the resolution of inflammation. The emergence of a TREM2^high^CD86^low^ macrophage signature is particularly notable. By downregulating CD86, a co-stimulatory molecule that drives inflammatory T-cell activation, while upregulating TREM2, a receptor central to debris recognition and clearance, these cells transition from pro-inflammatory sentinels to specialized effectors dedicated to tissue homeostasis and repair^59–62^. A parallel shift was observed in the neutrophil compartment, where rewilding downregulated CD86 and upregulated CD25, consistent with a less co-stimulatory and potentially immunomodulatory phenotype. Notably, rewilding also significantly increased the proportion of splenic CD4^+^CD25^+^ Tregs, pointing to broader engagement of IL-2/CD25-dependent regulatory circuits^15,63,64^. In this context, the neutrophil phenotype may reflect adaptation to a more regulatory immune milieu, potentially contributing to the containment of inflammatory amplification, although direct sequestration of IL-2 by neutrophils remains to be demonstrated. Higher CD44 expression on SSC^high^ neutrophils, concomitant to higher expression of CD25, further supports reprogramming of activation and trafficking states under rewilding conditions.

Another defining feature of the rewilded immune phenotype is the robust, multi-lineage upregulation of CD38 across plasmablasts, neutrophils, macrophages, and monocytes. As an ecto-enzyme, CD38 catalyzes the hydrolysis of extracellular NAD⁺, a damage-associated molecular pattern (DAMP) released during cellular stress and death^65–68^. Its broad induction across immune lineages therefore suggests enhanced capacity to sense and metabolically process extracellular danger cues, consistent with a reprogrammed state of immune readiness. Because CD38 can support both activation and immunomodulatory functions depending on cellular context, these findings are most consistent with coordinated remodeling of immune responsiveness. This rewilding induced shift, combined with the expansion and accelerated maturation of antibody-secreting cells toward more advanced differentiation states, underscores an immune system that is highly responsive yet tightly regulated. Concordantly, the selective rise in plasma IgA rather than a generalized increase in pro-inflammatory IgG isotypes reflects a shift in the host’s immune tone toward mucosal-associated immunity and active systemic regulation^5^.

The peripheral immune recalibration observed under rewilding has direct implications for CNS immunity. The immunological naïveté of SPF mice can blunt host responses to both pathogens and therapeutics, limiting the fidelity of preclinical models^6^. By establishing a mature, integrated host immune environment, our indoor rewilding approach provides a higher-fidelity platform that better reflects the cumulative immune history of the human host. This mature peripheral landscape, marked by enhanced metabolic surveillance, expanded regulatory T cells, and a diversified myeloid repertoire, likely constitutes an essential prerequisite for the microglial plasticity and transcriptional remodeling we observed within the brain^4,69^.

Within the CNS, rewilding did not simply suppress inflammation, it diversified and recalibrated it. The rewilded brain exhibited an expanded cytokine repertoire that integrated both pro-inflammatory (*IL-1β*, *IL-6*, *Ccl2*) and regulatory (*IL-10*) mediators, consistent with a transition from the monolithic, DAM-driven activation characteristic of SPF-5xFAD mice toward a mixed, context-dependent immune state^3,24,53^. This transition was strikingly mirrored at the protein level, where the systemic immune signature induced by rewilding (decreased CD86 alongside increased CD38 and CD25) was recapitulated in microglial populations in the brain. This molecular convergence between peripheral and central compartments suggests that rewilding coordinates a systemic-to-neuroimmune coupling that prevents microglia from becoming entrapped in a chronic pro-inflammatory state, instead sustaining a homeostatic surveillance capacity even in the presence of amyloid pathology^69^. By attenuating DAM-associated transcripts such as *Clec7a* and *Itgax* (CD11c) while simultaneously amplifying regulatory mediators, including *IL-10*, rewilding may recalibrate the neuroinflammatory response in 5xFAD mice toward a more adaptive phenotype^46^. Rather than simply dampening Aβ-associated inflammation, microbial and antigenic exposure appears to “train” innate immune pathways, promoting responsiveness while preventing uncontrolled neurotoxicity. This challenges the traditional binary view of neuroinflammation as purely detrimental and aligns with the growing understanding that microglial function is not fixed by genetics alone but is dynamically shaped by peripheral immune cues and ecological context.

Our transcriptomic analyses reveal that microbial and antigenic diversity exerts profound regulatory control over neuroimmune and metabolic gene expression in the brain. Rewilding redirected brain transcriptional profiles toward those observed in wild-caught mice, and, in the context of amyloid pathology, toward signatures found in human AD, reflecting both convergence and compensation between environmental and genetic determinants of disease. The co-upregulation of *Trem2*-*Clec7a* modules that define DAM, coupled with enhanced expression of canonical homeostatic markers such as *Gpr34*, supports the notion that ecological exposure promotes a state of microglial plasticity: one in which microglia acquire the priming signals necessary to respond to pathological challenges without permanently forfeiting their homeostatic surveillance identity, a balance frequently lost in sterile SPF environments^69–71^. Importantly, these transcriptional signatures were accompanied by the upregulation of mitochondrial and lysosomal pathways, indicating increased bioenergetic and phagocytic capacity, features central to microglial adaptation in neurodegeneration^72,73^.

Beyond microglial-specific transcriptional programs, rewilding re-engaged oxidative phosphorylation, complement activation, and antigen-presentation modules, overlapping with molecular signatures found in human AD brains. Notably, rewilded 5xFAD mice, but not their SPF counterparts, exhibited transcriptional profiles that more closely aligned with human AD hippocampus and entorhinal cortex datasets, as evidenced by increased DEG overlap and stronger cross-species pathway concordance. This region-specific convergence (absent in the postcentral gyrus and superior frontal gyrus) suggests that rewilding preferentially amplifies Aβ-relevant and regionally localized transcriptional programs rather than inducing a generalized inflammatory response. While we used human transcriptional signatures from females (to match our mouse-based studies), similar associations were seen in age-matched males (data not shown). Correspondingly, the human Alzheimer microglia (HAM) signature, largely absent under SPF conditions, was strongly induced in rewilded and wild mice, further underscoring the improved translational relevance of rewilded AD models. Together, these findings suggest that microbial and antigenic enrichment orchestrates a synchronized coupling between innate immune activation and metabolic adaptation, yielding a brain milieu that mirrors key molecular features of human AD while preserving dynamic regulatory control. Future studies integrating single-cell and spatial transcriptomics will be essential to delineate how environmental exposure reshapes immune-glial networks at the cellular level and to identify the circuits sustaining immune resilience.

Histological analyses further supported genotype-dependent remodeling of microglial morphology and Aβ engagement, broadly consistent with the transcriptional data. In WT brains, rewilding reduced the relative abundance of reactive microglia in both the cortex and hippocampus, supporting the idea that environmental exposure can reshape microglial states under basal conditions toward a less reactive profile. In 5xFAD brains, by contrast, rewilding had little impact on microglial state distribution, and microglia retained morphologies consistent with a reactive state, including enlarged somata and thickened, shortened processes, a hallmark of metabolic stress and inflammatory entrenchment^71,72^. This suggests that disease-associated microglial states may be comparatively resistant to short-term environmental modulation once pathology is established. Further, plaque-associated microglia were not numerically increased, yet showed significantly enhanced reactive microglial density per plaque, suggesting greater per-cell phagocytic engagement^74,75^. Whether this heightened physical interaction translates into net amyloid clearance remains an open question of temporal dynamics. Our 3-month rewilding protocol likely captures the re-education and recruitment phase of the neuroimmune response, in which microglia are functionally engaged but have not yet reached the threshold for significant plaque clearance. The increased microglial engagement observed without a concomitant reduction in plaque volume may alternatively reflect restoration of the “microglial barrier”, wherein microglia tightly encapsulate plaques to limit the diffusion of toxic Aβ oligomers toward surrounding neurites^74^. This model is consistent with the observed increase in mean plaque volume without a change in plaque count, suggesting modulation of plaque compaction dynamics rather than net accumulation. We therefore hypothesize that earlier-life exposure, together with more prolonged systemic immune priming mimicking chronic lifelong microbial exposure, may be required to convert this enhanced microglial engagement into measurable Aβ clearance.

The systemic-to-central immune interactions observed in this study point to a state of immunological resilience induced by rewilding, characterized by a dynamic equilibrium between activation and regulation. This contrasts sharply with the hyperreactive inflammatory milieu of SPF 5xFAD mice, where immune immaturity and exaggerated innate cascades may distort disease mechanisms and weaken translational fidelity. Such divergence likely contributes to the recurrent disconnect between preclinical outcomes and clinical reality, exemplified by the limited bench-to-bedside success of anti-Aβ immunotherapies and the emergence of amyloid-related imaging abnormalities (ARIA) in human trials. In humans, the neurovascular unit and blood-brain barrier operate within a mature immune milieu that is largely absent in SPF mice, potentially leading to systematic underestimation of vascular side effects associated with aggressive plaque clearance strategies^76–78^. These results highlight the need for AD models that better capture human immune complexity. While emerging systems such as cerebral organoids and iPSC-derived microglia xenografts offer powerful platforms for dissecting human-specific genetic mechanisms, they largely operate outside the organism-level context that shapes brain-immune interactions *in vivo*^79^. Our indoor rewilding framework is intended as a complementary, paradigm-level shift: rather than “humanizing” isolated cells in sterile settings, it reintroduces controlled ecological exposure to whole organisms to better align CNS pathology with a lifetime of systemic immune conditioning.

Future directions should focus on defining the causal microbial and antigenic drivers of brain immune plasticity and delineating how they interact with genetic AD risk factors such as *Apoe* and *Trem2*. Identifying the specific microbiome configurations, dietary components, and environmental signals that mediate neuroimmune reprogramming may enable precision rewilding strategies that enhance translational validity while maintaining experimental rigor. Additionally, exploring the roles of the circadian cycle, temperature, and other environmental variables^80^ may further refine the rewilding framework. Expanding this approach to other neurodegenerative pathologies, including Tau pathology, neurovascular dysfunction, and synucleinopathies, as well as to humanized AD risk models (e.g., APOE4, TREM2 variants) will be essential for determining the generalizability of rewilding-driven neuroimmune effects. In this way, environmental microbial complexity could become not merely a confounding variable to control, but a biological parameter to actively harness in the design of more predictive and ecologically grounded preclinical studies.

This study has several limitations. First, rewilded mice were exposed simultaneously to increased environmental complexity, greater physical activity, and a more diverse diet, each of which can independently influence AD-related outcomes. Although food intake was not quantitatively assessed, WT and 5xFAD mice were housed within the same enclosures under identical conditions, mitigating major environmental and dietary confounders between genotypes. Second, behavioral and cognitive performance were not directly evaluated in the current study, although our previous work with this rewilding platform^37^ demonstrated improvements in neuromotor and respiratory performance, and similar benefits might reasonably be expected here. Third, all animals were female, and sex-specific effects of rewilding on neuroimmune programming remain to be examined. Finally, the 3-month rewilding window, while sufficient to induce substantial immune and transcriptional remodeling, may not fully capture the long-term dynamics of microglial adaptation and amyloid clearance. Future investigations should systematically address how the duration of rewilding exposure, as well as age at onset, genetic background, and disease stage, modulate its neuroimmune effects.

In conclusion, this study provides the first comprehensive evidence that dirty mouse modeling reshapes neuroimmune and neuropathological processes within the CNS of a transgenic AD model. It establishes a foundation for exploring how ecological context intersects with the genetic and molecular determinants of AD, thereby bringing preclinical research closer to the real-world complexity of biological systems. Our findings extend the rewilding paradigm beyond systemic immunity, demonstrating that environmental microbial diversity reconfigures neuroimmune networks across multiple biological scales, from circulating lymphocytes to brain-resident microglia, and brings the transcriptional landscape of AD mice measurably closer to human disease. Ultimately, these insights may guide the development of more predictive, humane, and ecologically grounded experimental systems for studying and treating neurodegenerative diseases.

## MATERIALS AND METHODS

### Ethics statement

All procedures involving mice were carried out in accordance with the Canadian Council on Animal Care (CCAC) guidelines and were approved by the Animal Care Committee (CPAUL-2, approval number: 2024-1540). Female mice were chosen to reduce confounding factors related to breeding and limit aggression within the vivarium.

### Laboratory mice

Female 5xFAD transgenic mice (C57BL/6J genetic background; strain #034840-JAX; B6.Cg-Tg(APPSwFlLon,PSEN1*M146L*L286V)6799Vas/Mmjax) aged 6 weeks and their age-matched female wild-type (WT) littermates were purchased from MMRCC. After 2 weeks of housing under SPF laboratory conditions, the mice were weighed and randomly assigned to four experimental groups. The first two groups (SPF-WT and SPF-5xFAD) were kept under SPF conditions, while the other two groups (Rewild-WT and Rewild-5xFAD) were transferred to the indoor vivarium for rewilding. All groups were housed at ∼22°C and ∼45%RH under a 12:12-hour light/dark cycle, with food and water provided *ad libitum*. After three months, rewilded mice were recaptured using Havahart-type live traps. The vivarium environment, capture procedures, and post-capture handling were performed as previously described^37^. Finally, mice were weighed and euthanized concurrently with the SPF-housed mice.

### Wild-caught mice

Wild mice were captured in rural Québec City, Canada, using Longworth traps, which are specifically designed for the live capture of small mammals. Traps were set in the evening and left in place overnight at various strategic locations around a familial farm, including near garbage bins, at the entrances of buildings (the owner’s house, the sugar shack, the barn, the agricultural storage building, and the stable), and along structures where prior signs of murine activity had been observed. Traps were inspected early in the morning, and captured female mice were transported to the laboratory.

### Tissue collection and plasma preparation

All mice in this study were euthanized by decapitation without prior anesthesia, in accordance with institutional animal care and use guidelines and approved ethical protocols. Trunk blood was immediately collected into EDTA-coated tubes. For the N=8 cohort (see **Figure 1C**), a fraction of whole blood was kept fresh for flow cytometry, while the remainder was centrifuged (2000×g, 10 min, 4°C). The resulting plasma was aliquoted and stored at −80°C for subsequent MesoScale Discovery (MSD) assays (immunoglobulins and Aβ_40/42_). Brains were rapidly harvested and mid-sagittally hemisected. For the N=8 cohort, the right hemisphere and the spleen were processed fresh for cellular dissociation and flow cytometry. The left hemisphere was snap-frozen and cryo-milled on dry ice to be processed for qRT-PCR, Western blot, and MSD-based Aβ quantification (soluble and insoluble fractions of Aβ_38_, Aβ_40_, and Aβ_42_). For the transcriptomics and morphology cohort (N=6 cohort; see **Figure 1C**), the left hemisphere was processed for Nanopore sequencing, while the right hemisphere was fixed for immunohistochemistry (IHC) and morphological analysis.

### Flow cytometry

All samples were maintained on ice (in ice-cold 1xPBS for brain and spleen) until further processing. Single-cell suspensions were generated from each tissue and stained with fluorochrome-conjugated antibodies (**Supplementary Table 1**). Data were acquired using an Aurora spectral flow cytometer (Cytek Biosciences) and analyzed using FlowJo v10 software. Gating strategies were established based on fluorescence fluorescence-minus-one (FMO) controls and are presented in **Supplementary Figure 1A** and **1B**. Statistical analyses for flow cytometry were performed in R (RStudio 2026.01.0+392). Prior to modeling, outliers were systematically identified and excluded within each experimental group using a robust Z-score method based on the median absolute deviation (threshold ∣Z∣>3.5). To satisfy the assumptions of normality and homoscedasticity, cell abundance data were log(1+x) transformed, while population frequencies were logit-transformed. Median Fluorescence Intensity (MFI) values were normalized using an arcsinh transformation with a cofactor of 150. Statistical significance (defined at *p*<0.05) was assessed using linear models incorporating Genotype, Environment, and their interaction. Pre-specified simple contrasts were extracted using estimated marginal means (EMMeans). All p-values were derived from the global model’s pooled residual error to ensure a robust estimate of variance across all experimental groups. For selected panels, GraphPad Prism v10 was used to generate interaction plots and to fit two-way ANOVA full models for visualization of main effects and interactions (E, G, I reported as -log_10_(*p*) where indicated). Ratio metrics were analyzed after Log_10_-transformation, with raw ratio values displayed in figures where indicated.

### RNA extraction and quantitative RT-PCR

Around 80mg of cryo-milled brain was used for total RNA extraction with TRIzol Reagent (Life Technologies), according to the manufacturer’s instructions. RNA concentration and purity were assessed using a NanoQuant plate spectrophotometer (Infinite 2000, TECAN). Complementary DNA (cDNA) was synthesized from 2µg of total RNA using the High-Capacity cDNA Reverse Transcription Kit (Thermo Fisher Scientific, Cat# 4368814). qRT-PCR reactions were performed using 30 ng of cDNA using PowerTrack SYBR Green Master Mix (Thermo Fisher Scientific, Cat#A46109) on a LightCycler 480 II system (Roche). Primer sequences (1 µM each) are listed in (**Supplementary Table 2**), and their specificity was verified using NCBI Primer-BLAST. The amplification protocol consisted of the following steps: 50°C 2min, 95°C 10min, [95°C 30sec, 60°C 30sec] x40, 95°C 1min, 55°C 30sec, 95°C 30sec. Relative gene expression levels were determined using the 2^−ΔΔCt^ method, using endogenous Gapdh as a normalization gene. Data were log_2_-transformed prior to statistical analysis using GraphPad prism v10.

### Nanopore RNA Sequencing

Total RNA (2µg per sample) was extracted from the left brain hemisphere as described above and sent to the Integrated Genomics Meta-Platform at CHU Sainte-Justine (Montréal, QC, Canada) for bulk (whole tissue) sequencing. Libraries were prepared from total extracted RNA using the cDNA-PCR sequencing kit (SQK-PCB114-24) from Oxford Nanopore. The samples were loaded onto 3 FLO-PRO114M flow cells and sequenced PromethION 24 at the CRA-CHUSJ Integrated Genomics Meta-Platform (MPGI) for 72 hours. Pod5 files were base called with Dorado v.0.8.0 (Oxford Nanopore Technologies) using model sup@v5.0.0. Reads with a Phred Q score of 10 or more were kept for downstream analysis. Transcripts quantification and alignment were done using oarfish v.0.6.5 ^81^ and Mus musculus GRCm39 reference transcriptome with parameters ‘--seq-tech ont-cdna’, ‘--filter-group no-filters’ and ‘--model-coverage’. Differential expression analysis was done using the R package DESeq2 v.1.48.2 ^82^ and counts were normalized with the default method, median of ratios.

### Protein extraction and immunoblotting

Cryo-milled brain was used to isolate soluble and insoluble protein fractions. Specifically, 100 mg of frozen brain powder were lysed in RIPA buffer composed of 50 mM Tris-HCl (pH 8), 150 mM NaCl, 1% Nonidet P-40, 0.5% sodium deoxycholate, 0.1% SDS, 1 mM phosphatase inhibitors, 1 mM PMSF, and one Complete Mini EDTA-free protease inhibitor tablet (Roche Life Science). Lysates were centrifuged at 100,000×g for 20 min at 4 °C, and the supernatant containing the soluble fraction was collected. The remaining pellet, representing the insoluble fraction, was resuspended in 70% formic acid and centrifuged at 25,000×g for 20 min at 4 °C. The resulting supernatant was evaporated under a fume hood for three days, and the dried residue was resuspended in 5M guanidine hydrochloride diluted in 0.05M Tris-HCl (pH 8). Protein concentrations were quantified using the Pierce BCA Protein Assay Kit (ThermoFisher Scientific). For immunoblotting, 20μg of protein per sample were resolved on 15% SDS-PAGE gels and transferred onto nitrocellulose membranes (Bio-Rad). Membranes were blocked for 1 h at room temperature with 5% non-fat dry milk in TBS containing 0.1% Tween-20, followed by overnight incubation at 4 °C with the appropriate primary antibodies (listed in **Supplementary Table 3**). After washing, membranes were incubated with HRP-conjugated secondary antibodies and developed using the Immobilon® ECL UltraPlus Western HRP Substrate (Millipore). Total protein staining (Ponceau S) was used for normalization. Signals were visualized using a ChemiDoc Imaging System (Bio-Rad), and densitometric quantification was performed with ImageJ software (version 1.53). Statistical analysis used GraphPad prism v10.

### Multiplex MSD quantification of Amyloid beta and immunoglobulins

The quantification of human amyloid beta peptides, Aβ_38_, Aβ_40_, and Aβ_42_, in the brain was performed using the V-PLEX Aβ Peptide Panel 1 kit 6E10 (#K15200E, Meso Scale Discovery, Maryland, USA). Whereas the quantification of human amyloid beta peptides, Aβ_40_ and Aβ_42_, in the plasma was performed using the R-PLEX 4G8 kit (#L45SA-1; #F209M-3; #K1509MR-2; #K1509LR-2; #F209L-3). The quantification of plasma immunoglobulins was performed using the T-PLEX Mouse Isotyping Panel 1 (IgA, IgG1, IgG2a/c, IgG2b, IgG3, IgM) (#K15183B). All procedures were according to the manufacturer’s instructions. Plates were read on a MESO^®^ QuickPlex SQ-120 Multiplex imager. Data were analyzed using MSD-Workbench software. Results were log_10_-transformed prior to statistical analysis to account for skewed distributions, using GraphPad prism v10.

### Immunofluorescence

The right brain hemispheres (N=6 per group) were fixed in 4% paraformaldehyde (PFA) for 12h and then transferred to PBS-sucrose 30% for 24h for cryopreservation. The half-brains were then transferred to isopentane at −40°C on dry ice for 10s for rapid freezing. The half brains were kept at −80°C until cold sectioning with a cryostat microtome (thickness, 16 mm). The resulting cryosections were preserved from contamination in PBS-Azide 0.2% until immunostaining. Cryosections were rinsed three times in PBS, followed by successive immersions in 70% and 80% ethanol, Thioflavin S (T1892-25G, Sigma Aldrich, USA) was prepared at 1% in 80% ethanol, and the solution was syringe-filtered to remove any undissolved particles. Slides were then incubated in the Thioflavin S solution for 15 minutes at room temperature under agitation, protected from light. After incubation, slides were rinsed sequentially in 80% and 70% ethanol followed by two washes in PBS. Sections were subsequently permeabilized with PBS-Triton 0.2% and saturated with Normal Goat Serum 1/100 (Vector Laboratories; Cat. No. S-1000) for one hour at room temperature and then labeled with primary antibodies anti-Iba1 1/500 (Rabbit, #019-19741, Wako). Afterwards, sections were incubated with secondary antibodies goat anti-rabbit 568 1/500 (#A11036, Invitrogen) for 1 hour at RT. The sections were then incubated with DAPI (D1306, Invitrogen) and bathed in 70% ethanol and Sudan black (Sigma, Canada) to eliminate autofluorescence. The sections were then mounted on Superfrost slides (ThermoFisher) using Fluoromount-GTM Mounting Medium (Invitrogen, 00495802). The slides were finally digitized using a slide scanner (Axioscan Z1, Zeiss).

### Image analysis

High-resolution images obtained from the slide scanner were analyzed using a flexible algorithm developed by the imaging platform of our research center under MathWorks MATLAB 2018b software. Briefly, the code includes a multi-CPU-GPU batch image processor that reads raw images, microscope metadata, and experimental factors to serve 4D matrices to a custom plugin written to detect 3D objects labeled in red (microglia) and green (Aβ plaques). This automated pipeline quantified, within the cortex (C) and the hippocampus (HX), the object properties such the size, the number, and the fluorescence intensities. The property values were aggregated along samples and groups to produce tissue measurements in terms of means ± standard error of the mean (SEM). Object intersections were determined from voxel lists to get microglia-plaque colocalization. The plugin further distinguished distinct microglial morphologies based on soma size and ramification complexity (Homeostatic, Reactive, Amoeboid), providing an integrated assessment of microglial activation states. All quantitative measures were normalized to the analyzed mask volume. Group comparisons were performed using an ANOVA 2-way statistical test at 5% *p*-value threshold followed by a Holm-Bonferroni post-hoc test under OriginLab (OriginPro 2015 software).

### Patient and public involvement

Patients and/or the public were not involved in the design, conduct, reporting, or dissemination plans of this research.

## Acknowledgements

We thank all family members of *La ferme du vieux caveau* farm located in Saint-Joachim, Québec, for generously providing soil and other organic materials for our indoor enclosures and for allowing us to capture Wild mice. We also thank Sébastien Poulin and all staff members of the IUCPQ animal facility for their support and guidance. Our gratitude extends further to the CRCHUQ animal facility team for their continuous assistance. Finally, we thank Vincent Desrosiers from the cytometry platform at the CHU de Québec-Université Laval Research Center for guidance and data analysis.

## CRediT authorship contribution statement

**Mohamed Lala Bouali**: conceptualization, methodology, investigation, formal analysis, visualization, data curation, writing - original draft, writing - review & editing of the final manuscript. **Amir Mohamed Kezai**: conceptualization, methodology, investigation, formal analysis, writing - original draft. **Marc Bazin:** formal analysis. **Papa Yaya Badiane**: investigation. **Nasim Eskandari**: investigation. **Véronique Lévesque**: investigation. **Justin Robillard**: resources. **Steeve D. Côté**: resources. **Denis Soulet**: resources. **Cyntia Tremblay**: investigation. **Frédéric Calon**: resources. **Françoise Morin**: investigation, formal analysis. **Luc Vallières**: resources, formal analysis, validation, writing - review & editing of the final manuscript. **Sébastien S. Hébert**: funding acquisition, conceptualization, project administration, resources, supervision, validation, data curation, formal analysis, visualization, writing - review & editing of the final manuscript. All the authors have read and approved the final manuscript.

## Materials & Correspondence

The authors confirm that the datasets, including raw data, supporting the findings of the current study, are available from the corresponding author. The raw Nanopore FASTQ files were unexpectedly lost during manuscript preparation due to circumstances beyond the authors’ control. Processed data, including normalized gene-expression matrices, are provided with this manuscript and remain fully available for review. All corresponding tissue specimens and extracted total RNA are preserved and can be used to regenerate sequencing datasets upon reasonable request to the corresponding author.

