## Supplementary Figures and Tables for "Natural Microbial Enrichment Modulates Microglial States and Transcriptional Programs Relevant to Alzheimer’s Disease"

Supplementary Material

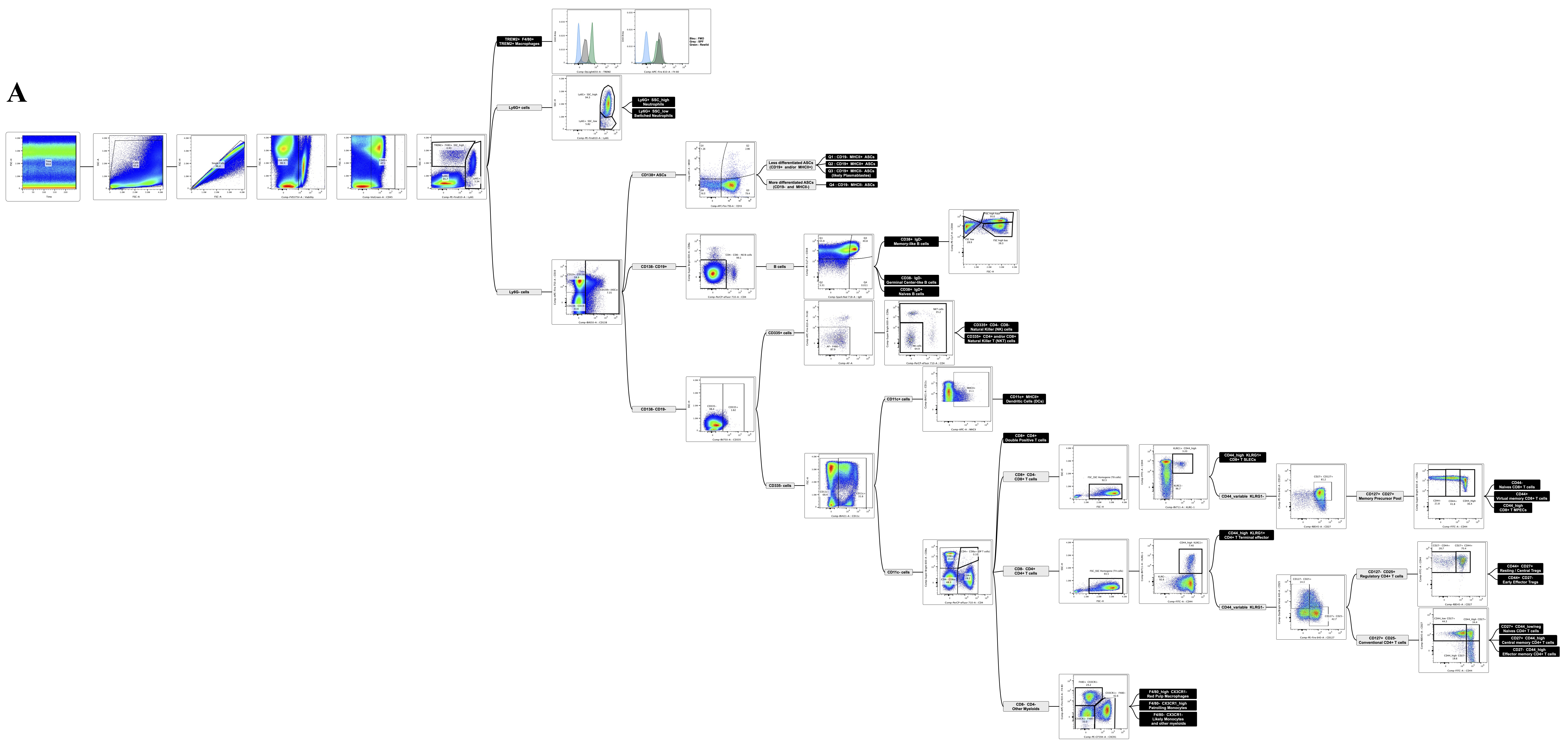

**Supplementary Figure 1. Representative gating strategy for spleen and blood samples.** Sequential gating was performed starting from time, FSC/SSC, singlets, live cells, and CD45^+^ leukocytes, followed by specific lineage markers. A representative Spleen **(A)** and Blood **(B)** sample from a same Rewild-5xFAD mouse is shown. See the online version for an enlarged view.

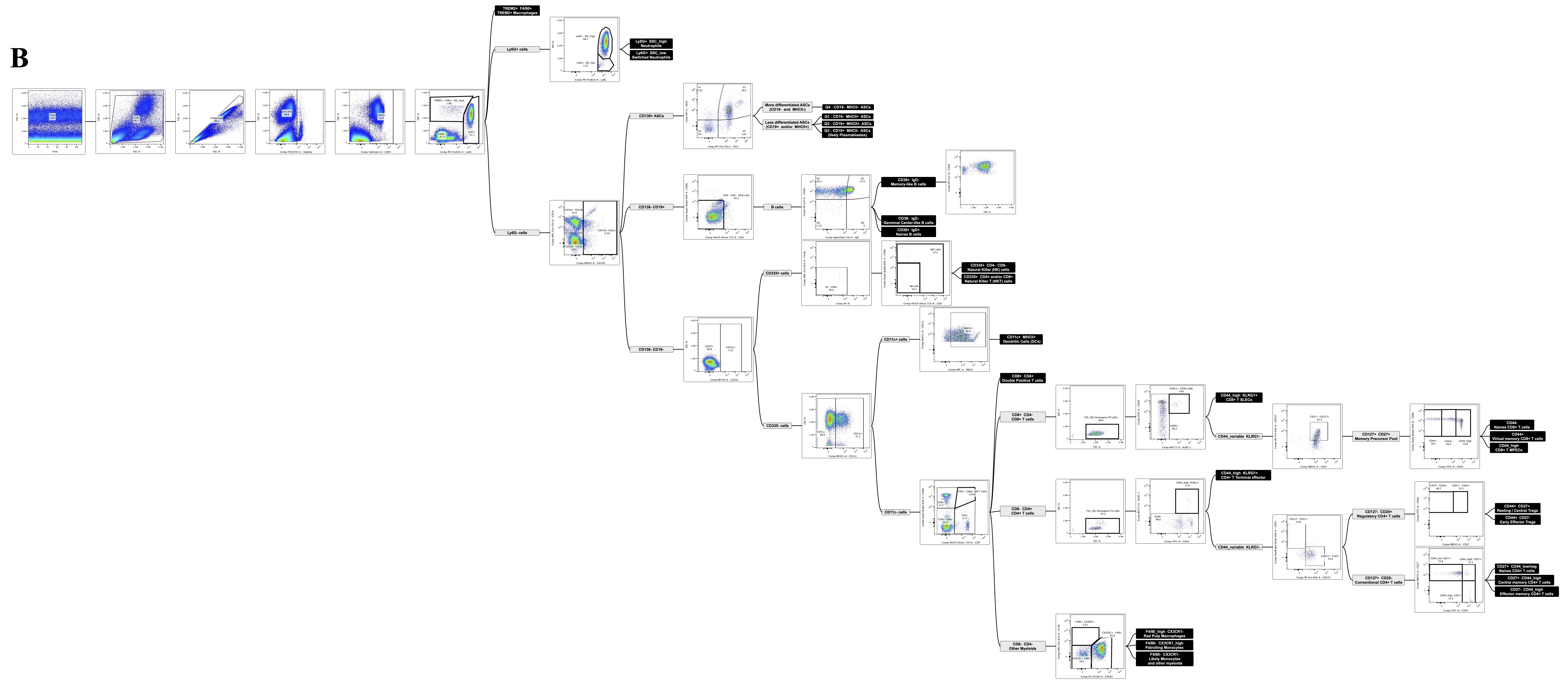

**Supplementary Figure 1. Representative gating strategy for spleen and blood samples.** Sequential gating was performed starting from time, FSC/SSC, singlets, live cells, and CD45^+^ leukocytes, followed by specific lineage markers. A representative Spleen **(A)** and Blood **(B)** sample from a same Rewild-5xFAD mouse is shown. See the online version for an enlarged view.

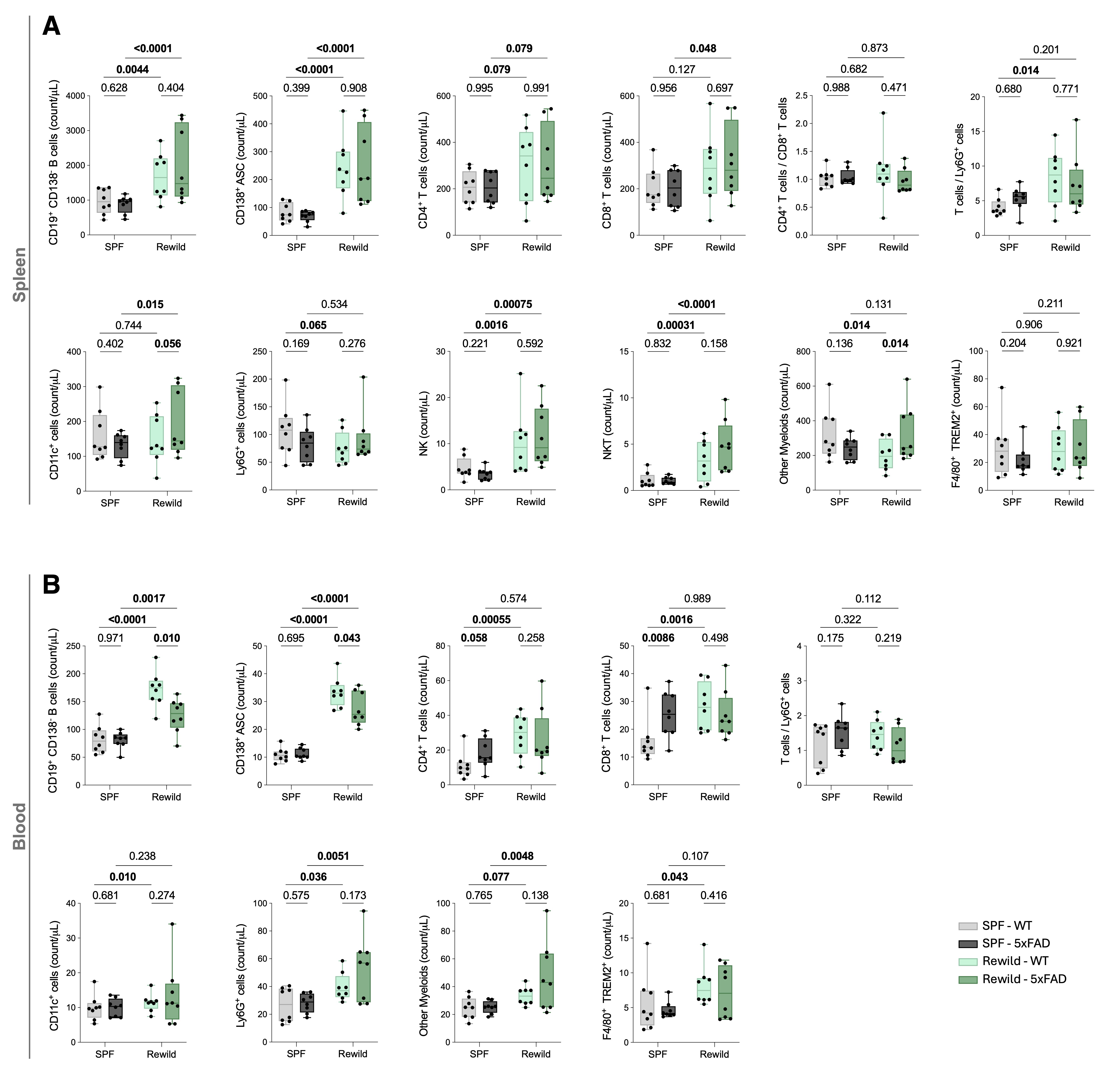

**Supplementary Figure 2.** **Count of leukocyte subsets in spleen and blood.** Flow cytometry-based volumetric abundances (counts/µL of acquired sample; acquisition target ~100 µL per sample) of major leukocyte populations measured in spleen **(A)** and blood **(B)** from female mice housed under SPF conditions or in the indoor vivarium (Rewild): SPF-WT, SPF-5xFAD, Rewild-WT, and Rewild-5xFAD (N = 8 per group). The Total T-cells/Ly6G⁺ cells ratio and the CD4^+^/ CD8^+^ T-cells ratio are included where indicated. Statistical analyses and outlier handling were performed as described in Methods. Ratios were analyzed after log_10_ transformation, whereas raw ratio values are displayed.

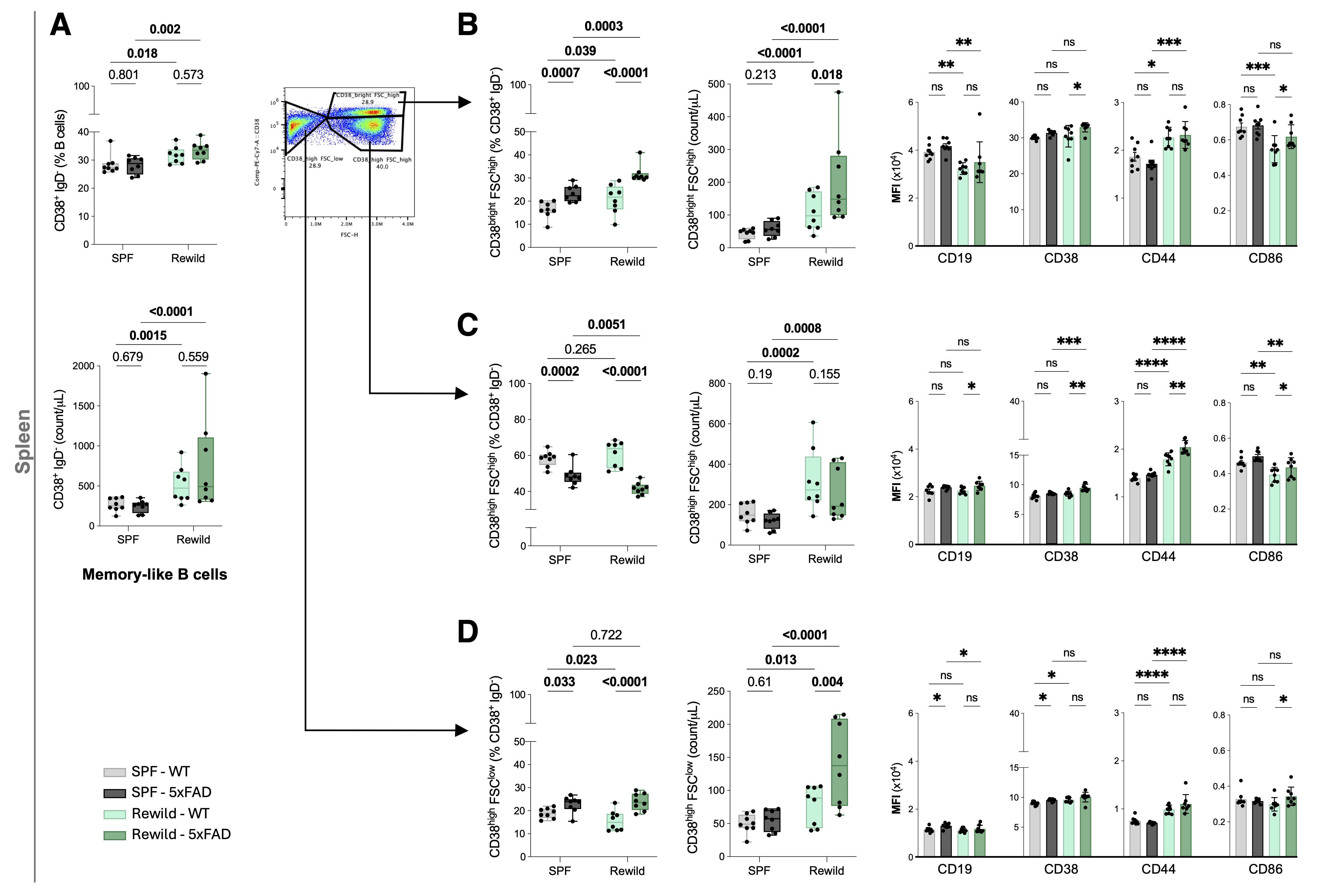

**Supplementary Figure 3.** **CD38/FSC-defined heterogeneity within splenic memory-like B cells.** Flow cytometry analysis of splenic memory-like B cells from female SPF-WT, SPF-5xFAD, Rewild-WT, and Rewild-5xFAD mice (N = 8 per group). Memory-like B cells were operationally defined as CD38⁺IgD⁻ (as in **Figure 3**) and were further partitioned based on CD38 intensity and cell size (FSC-H). **(A)** Total CD38⁺IgD⁻ memory-like B cells in spleen shown as numbers (counts/µL) and as proportions of B lymphocytes, with representative gating illustrating the CD38/FSC-based subdivision quantified in panels B-D. **(B-D)** Quantification of CD38/FSC-defined subfractions within the memory-like compartment: CD38^bright^FSC^high^ **(B)**, CD38^high^FSC^high^ **(C)**, and CD38^high^FSC^low^ **(D)**. For each subfraction, results are shown as numbers (counts/µL) and as proportions of CD38⁺IgD⁻ cells, alongside marker expression (MFI) for CD19, CD38, CD44, and CD86. Statistical analyses and outlier handling were performed using the pipeline described in Methods. **p* < 0.05, ***p* < 0.01, ****p* < 0.001, *****p* < 0.0001; ns: not significant.

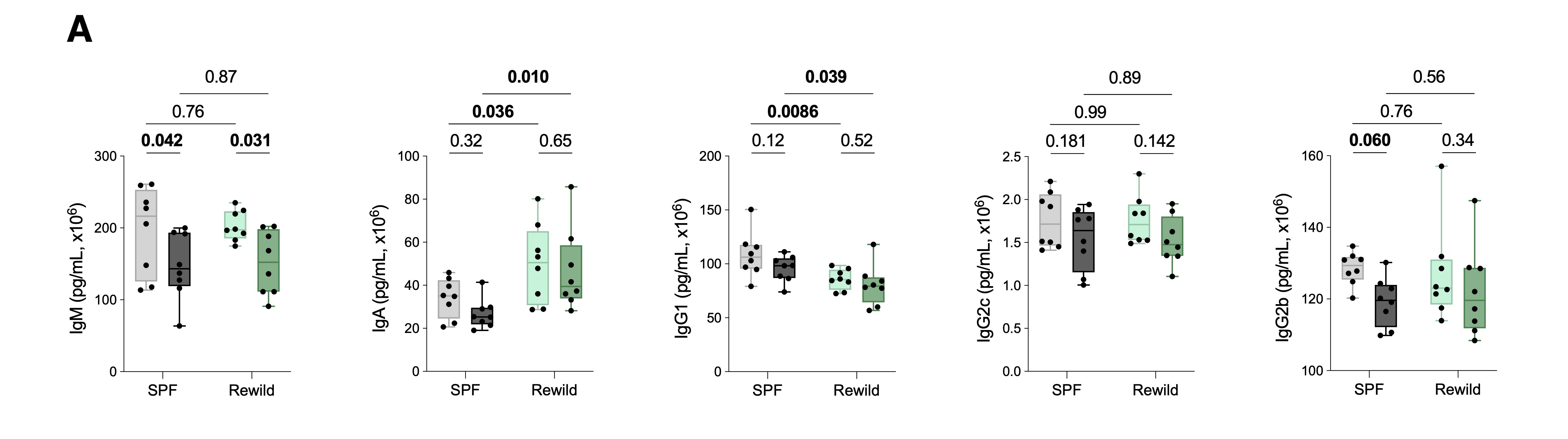

**Supplementary Figure 4.** **Plasma immunoglobulins profiling.** Plasma immunoglobulin concentrations (IgM, IgA, IgG1, IgG2b, IgG2c) were quantified by Meso Scale Discovery (MSD) immunoassays in female SPF-WT, SPF-5xFAD, Rewild-WT, and Rewild-5xFAD mice (N = 8; flow cytometry cohort). Values are reported as pg/mL (x10^6^) as indicated. Statistical testing was performed on log_10_-transformed data using a two-way ANOVA.

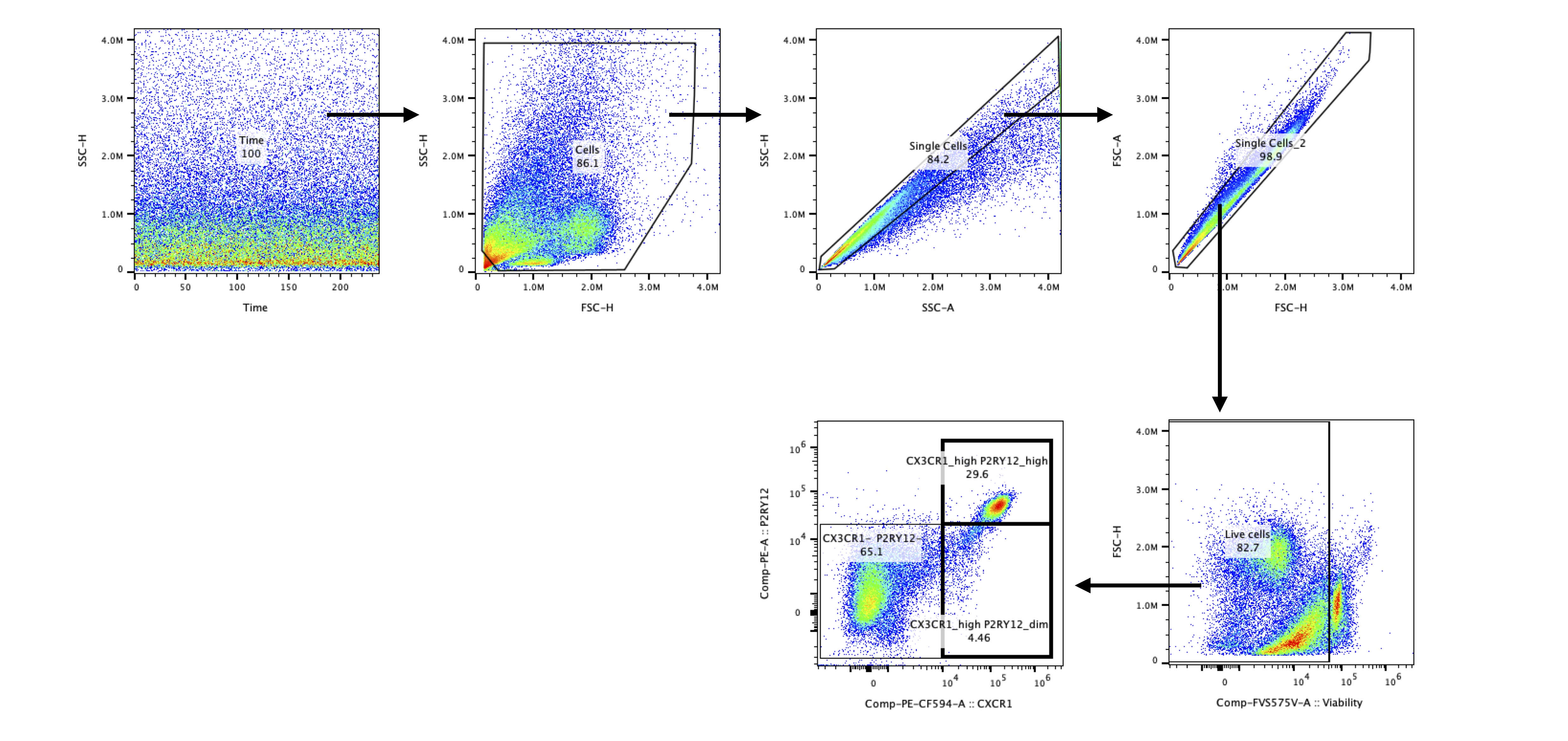

**Supplementary Figure 5.** **Representative gating strategy for brain samples.** Sequential gating was performed starting from time, FSC/SSC, singlets, live cells, then identification of microglia via P2RY12 vs CX3CR1 plot.

**Supplementary Table 1.** List of fluorochrome-conjugated antibodies and reagents used in flow cytometry for blood, spleen and brain leukocyte profiling.

| Antigen / Reagent | Fluorochrome | Supplier | # Catalog |
| --- | --- | --- | --- |
| Fixable Viability Stain | FVS575V | BD Biosciences | 565694 |
| TREM2 | DyLight 650 | Novus | FAB17291W |
| P2RY12 | PE | BioLegend | 848004 |
| MHCII | APC | Invitrogen | APC-65122-100UG |
| Ly6g | PE-Fire 810 | BioLegend | 127673 |
| KLRG1 | Brilliant Violet 711 | BD Biosciences | 564014 |
| IgD | AlexaFluore700 | BioLegend | 405730 |
| Fc block | - | BioLegend | 103116 |
| F4/80 | APC-Fire 810 | BioLegend | 123166 |
| CX3CR1 | PE-CF 594 | BD Biosciences | 567823 |
| CD8a | Super Bright 600 | Invitrogen | 63-0081-82 |
| CD86 | VioBright 515 | Miltenyi Biotec | 130-122-136 |
| CD45 | VioGreen | Miltenyi Biotec | 130-123-900 |
| CD44 | FITC | Invitrogen | 11-0441-82 |
| CD4 | PerCP-eFluor 710 | Invitrogen | 46-0041-82 |
| CD38 | BrilliantViolet 480 | Invitrogen | 414-0381-82 |
| CD335 | Brilliant Violet 750 | BD Biosciences | 746875 |
| CD27 | RB545 | BD Biosciences | 756643 |
| CD25 | StarBright Violet 440 | Bio-Rad | MCA1260SBV440 |
| CD19 | APC-Fire 750 | BioLegend | 115558 |
| CD138 | Brilliant Violet 650 | BioLegend | 142518 |
| CD127 | PE-Fire 640 | BioLegend | 158214 |
| CD11c | BV421 | Invitrogen | 404-0114-82 |
| CD11b | StarBright Blue 700 | Bio-Rad | MCA711SBB700 |

**Supplementary Table 2.** List of forward and reverse primers designed for amplification of murine cytokine, chemokine and housekeeping genes. Primer sequences were used to quantify mRNA expression in brain tissue by SYBR Green-based qRT-PCR.

|  | Forward | Reverse |
| --- | --- | --- |
| Target murine genes | | |
| Tnf | TGCCTATGTCTCAGCCTCTTC | GAGGCCATTTGGGAACTTCT |
| Clec7a | TGCCTATGTCTCAGCCTCTTC | GAGGCCATTTGGGAACTTCT |
| IL-1a | ATGGTTCTGGGAGGATGGAT | GCTTTCCTGGGGAGCTGTAT |
| IL-1b | TTGGTTAAATGACCTGCAACA | GAGCGCTCACGAACAGTTG |
| Nlrp3 | ATCTTTTGGGGTCCGTCAACT | GCAACTGTTCCTGAACTCAACT |
| IL-6 | ATTACCCGCCCGAGAAAGG | CATGAGTGTGGCTAGATCCAAG |
| Ccl2 | ACCACGGCCTTCCCTACTTC | TCCACGATTTCCCAGAGAACA |
| Itgax | GCTACAAGAGGATCACCAGCAG | GTCTGGACCCATTCCTTCTTGG |
| IL-10 | ATGGAGCCTCAAGACAGGAC | GGATCTGGGATGCTGAAATC |
| Housekeeping murine gene | | |
| Gapdh | ACAAAATGGTGAAGGTCGGTG | TGGCAACAATCTCCACTTTGC |

**Supplementary Table 3.** List of primary and secondary antibodies used for protein detection by immunoblot. Dilutions used, host species, suppliers, and catalog numbers are indicated.

| Protein | Dilution | | Species | Supplier | # Catalog |
| --- | --- | --- | --- | --- | --- |
| Primary antibodies | | | | | |
| Iba1 | | 1/1000 | Mouse monoclonal | Invitrogen | MA5-27726 |
| Gfap | | 1/1000 | Mouse monoclonal | BioLegends | 835301 |
| Secondary antibodies | | | | | |
| HRP-conjugated IgG (HC+LC) | | 1/5000 | Goat anti-mouse | Jackson laboratories | 115-035-003 |
